# Partial reprogramming reveals lineage-specific uncoupling of rejuvenation, function, and recoverability in human cells

**DOI:** 10.64898/2026.11.09.764947

**Authors:** Kaho Washizu, Xun Wang, Julia M. Draper, Shinya Yamanaka, Akira Kunitomi

## Abstract

Partial reprogramming is therapeutically useful only if cells keep its benefits while recovering their lineage identity and function after the reprogramming factors are withdrawn. Whether the relationship between reprogramming depth and recovery differs between lineages is unknown. Using reversible, traceable Sendai-OSKM, we prospectively isolated intermediate reprogramming states from primary human fibroblasts, cytotoxic T cells, and aortic endothelial cells and followed their recovery. Outcomes diverged: fibroblasts kept a reduced DNA methylation age but retained stromal remodeling and incomplete proliferative recovery; T cells recovered only from shallow reprogramming, with enhanced effector function; endothelial cells rapidly recovered identity, kept a reduced methylation age, and markedly improved vascular function. None of this tracked iPSC derivation efficiency. Multiome and knockdown analyses identified TOX as a shared progression dependency, whereas its candidate downstream programs were predominantly lineage-specific. Partial reprogramming involves a lineage-specific tradeoff between rejuvenation and recovery that must be optimized one lineage at a time.

## Introduction

Age-associated molecular and epigenetic alterations progressively compromise tissue function^1–6^. However, some age-associated alterations are reversible in the lab, which has motivated efforts to repair aging tissues by resetting cellular states without replacing the cells themselves. Complete reprogramming with OCT4, SOX2, KLF4, and MYC (OSKM) can erase age-associated molecular features^7–9^. However, it entails the acquisition of pluripotency and the loss of somatic identity, thereby raising substantial safety concerns for in vivo applications, particularly the risk of teratoma formation^10^. Several alternatives to complete reprogramming have therefore been investigated. Transient reprogramming has improved age-associated molecular and functional phenotypes in vivo^11,12^ and in cultured human cells^13,14^, while reversible reprogramming has promoted regenerative plasticity^15^. Several modes of transient reprogramming have been investigated, including brief exposure to reprogramming factors of dermal fibroblasts and endothelial cells derived from iliac arteries and veins, which did not lead to overt identity loss^13^, and progressively longer OSKM exposure followed by extended factor-free recovery in human dermal fibroblasts, which revealed identity reacquisition and an exposure-duration-dependent optimum for molecular rejuvenation^14^. One study mapped these dynamics at single-cell resolution in murine adipogenic and mesenchymal stromal cells, and found that partial reprogramming restored youthful gene expression while suppressing somatic identity only transiently, and that the two effects were not tightly coupled^16^. Another found that a single fixed OSK exposure attenuated endothelial-to-mesenchymal transition and cellular senescence in human aortic endothelial cells and lowered blood pressure in hypertensive mice^17^. Nevertheless, no study has systematically compared multiple primary human lineages to determine how reprogramming depth affects the cells’ ability to retain the benefits of reprogramming and to recover their lineage identity and function after withdrawal of the reprogramming factors. This information is necessary to define, for each cell type, a partial reprogramming window that yields the desired benefits without compromising post-withdrawal recovery.

In addition, the endogenous processes linking OSKM-induced progression to intermediate reprogramming states and post-withdrawal outcomes remain poorly defined. The ability of OSKM to induce pluripotency across diverse somatic cell types suggests that OSKM engages common core mechanisms. Yet reprogramming efficiency and trajectories vary markedly with cell of origin, and cross-lineage studies have identified both shared late-stage features and cell-type-specific sequences of identity loss and pluripotency acquisition^18,19^. These observations raise the question of whether partial reprogramming proceeds through equivalent intermediate states across lineages or instead combines shared regulatory dependencies with lineage-specific molecular programs.

Here, using a traceable, reversible Sendai-OSKM system, we prospectively isolated graded intermediate-state populations from primary human fibroblasts, cytotoxic T cells, and aortic endothelial cells and followed their recovery after OSKM withdrawal. Our findings argue against defining partial reprogramming by the duration of factor exposure or inferring the cells’ ability to recover from the efficiency with which they can be fully reprogrammed into iPSCs. Interventions intended to modify age-associated phenotypes should therefore be judged not by molecular-age changes alone, but by whether cells recover their lineage identity and specialized function after factor withdrawal. Our findings also identify the HMG-box factor TOX as a shared reprogramming dependency that coexists with lineage-specific intermediate states and recovery responses. TOX provides a common mechanistic entry point from which we can ultimately dissect the origins of lineage-divergent mechanisms and outcomes of partial reprogramming.

## Results

### OSKM induces gradual reprogramming with different kinetics in three human lineages

Studying partial reprogramming requires a system in which exogenous OSKM expression can be both controlled and traced, so that cells which actually received OSKM can be identified while alive. Because diverse somatic cell types differ widely in their permissiveness to gene transfer, we also needed a vector with high delivery efficiency. We therefore adapted our previously reported, improved temperature-sensitive Sendai virus (SeV) reprogramming platform^20^ by incorporating GFP into SeV-KOS/TS12 and OFP into SeV-cMYC/TS15 (Figure 1A). Although SeV-KOS/TS12 already encodes KLF4, we retained SeV-KLF4/TS12 as a third vector because, in preliminary optimization of this system, its inclusion increased iPSC colony formation by more than an order of magnitude relative to the two-vector combination. Sorting cells positive for both GFP and OFP enriched downstream analyses for cells receiving all four OSKM factors. As a control, cells were infected in parallel with matched temperature-sensitive fluorescent SeV control vectors carrying GFP and OFP but lacking the OSKM cassettes, together with an empty SeV vector in place of SeV-KLF4/TS12. This matched control-vector condition is referred to as Ctrl throughout. Flow cytometry showed that GFP+OFP+ cells receiving the matched control vectors retained the marker profile of the starting somatic cells in all three cell types (Figure S1B).

**Figure 1.**
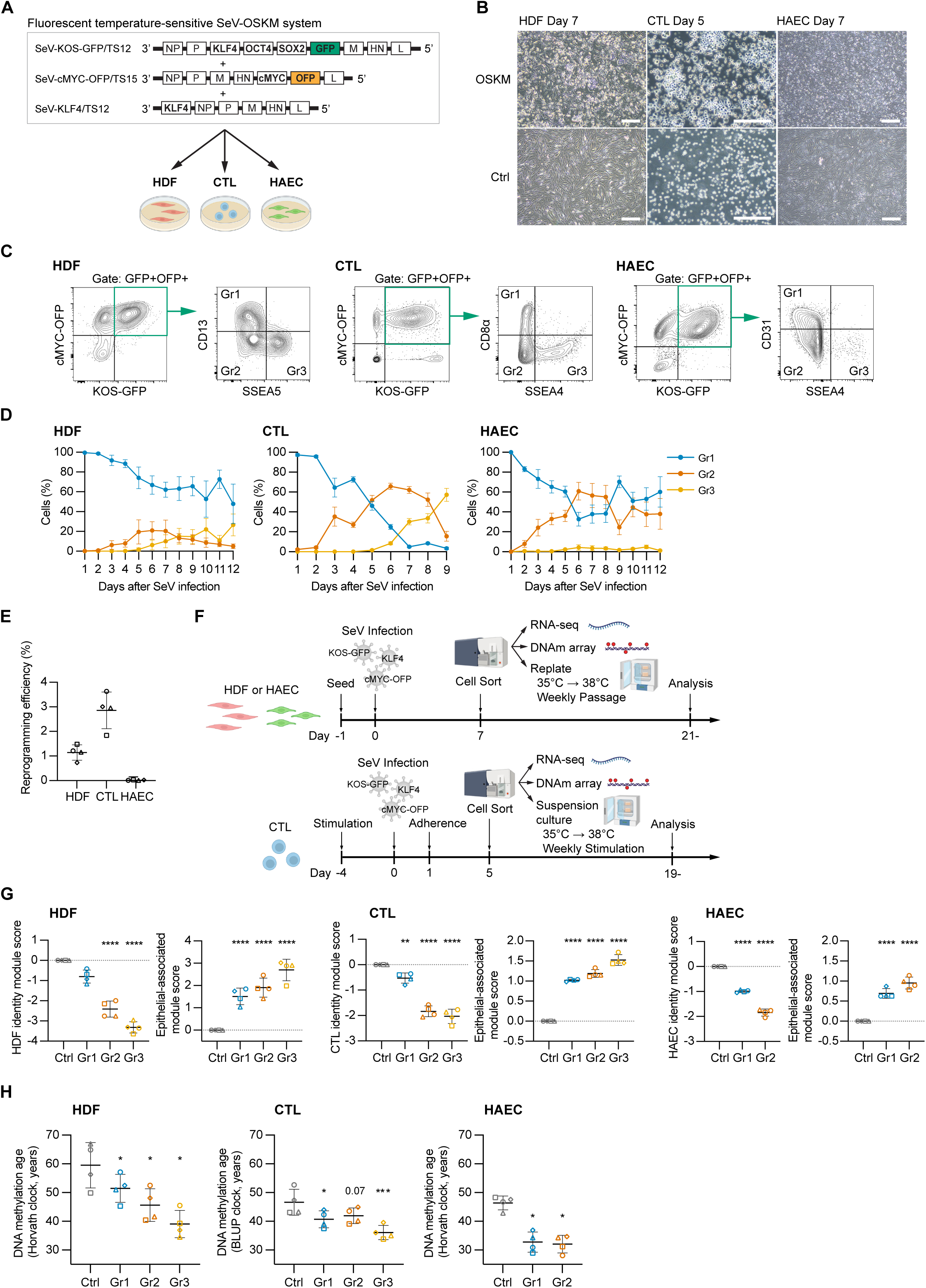
A fluorescent temperature-sensitive SeV-OSKM system resolves graded, lineage-specific reprogramming trajectories across three human somatic cell types. (A) Schematic of the fluorescent temperature-sensitive SeV-OSKM system used across three human somatic cell types. GFP was inserted into SeV-KOS/TS12 and OFP into SeV-cMYC/TS15, enabling live enrichment for cells receiving both fluorescent components of the OSKM vector set. All three vectors, including SeV-KLF4/TS12, were temperature-sensitive. Box lengths are not to scale. (B) Representative microscopy images after SeV-OSKM infection in HDFs, CTLs, and HAECs, showing cell type-specific morphological responses to SeV-OSKM expression. Scale bars, 200 µm. (C) Flow cytometry strategy used to stratify GFP+OFP+ cells into graded partial reprogramming groups. HDFs were subdivided by CD13 and SSEA5, CTLs by CD8α and SSEA4, and HAECs by CD31 and SSEA4 to define Gr1, Gr2, and Gr3 along the partial reprogramming continuum. (D) Time-course quantification of Gr1, Gr2, and Gr3 fractions among GFP+OFP+ cells after SeV-OSKM infection in HDFs, CTLs, and HAECs. Marker-defined partial reprogramming states dynamically redistributed after infection, revealing rapid progression in CTLs, more gradual progression in HDFs, and limited Gr3 accumulation in HAECs. Dots represent means from four independent donors; error bars indicate SD. (E) iPSC reprogramming efficiency in HDFs, CTLs, and HAECs after SeV-OSKM infection. The central horizontal line indicates the mean of four independent donors, and error bars indicate SD. Open symbols indicate individual donors: HDFs, circle = 54F, square = 48F, triangle = 56M, diamond = 48M; CTLs, circle = 58M, square = 55M, triangle = 56F, diamond = 54F; HAECs, circle = 63M, square = 61M, triangle = 51F, diamond = 50F. (F) Experimental workflow for sorting graded partial reprogramming states and collecting bulk RNA-seq and DNA methylation array samples. HDFs and HAECs were sorted on day 7, whereas CTLs were sorted on day 5. After sorting, cultures were shifted from 35°C to 38°C to suppress replication of the temperature-sensitive SeV genome and promote loss of SeV-OSKM expression. (G) Cell type-specific somatic identity module scores and epithelial-associated module scores derived from MET-associated genes at the sorting time point. In HDFs, the latter captures canonical MET; in CTLs and HAECs, it denotes acquisition of the same epithelial-associated gene set rather than a literal mesenchymal-to-epithelial transition. For visualization, donor-level module scores are shown as differences from the matched Ctrl score; statistical comparisons and displayed significance values were calculated from the corresponding original (uncentered) donor-level scores. The central horizontal line indicates the mean of four independent donors, and error bars indicate SD; open symbols indicate the same individual donors listed in panel E. For comparisons among more than two related conditions, repeated-measures one-way ANOVA followed by Dunnett’s multiple-comparisons test versus Ctrl was used. \**p* < 0.05, \*\**p* < 0.01, \*\*\**p* < 0.001, \*\*\*\**p* < 0.0001. (H) DNA methylation age estimates at the sorting time point. HDF and HAEC ages were estimated using the Horvath clock. For CTLs, BLUP estimates are shown as the primary readout; Horvath and IntrinClock estimates showed greater variability in this dataset and are shown in Figures S1E and S1F. The central horizontal line indicates the mean of four independent donors, and error bars indicate SD; open symbols indicate the same individual donors listed in panel E. For comparisons among more than two related conditions, repeated-measures one-way ANOVA followed by Dunnett’s multiple comparisons test versus Ctrl was used. \**p* < 0.05, \*\*\**p* < 0.001; exact *p* values are shown where indicated.

We applied this system to human dermal fibroblasts (HDFs), cytotoxic T cells (CTLs), and human aortic endothelial cells (HAECs), using four donors per cell type; donors ranged in age from 48 to 63 years, and both sexes were represented within each cell type. These lineages were chosen with complementary rationales. HDFs are the cell type used in most reprogramming and partial-reprogramming studies and therefore provide a benchmark against which state-resolved measurements can be compared. CTLs are directly relevant to adoptive cell therapy, and full reprogramming followed by redifferentiation can rejuvenate antigen-specific T cells^21,22^. We were therefore curious whether partial reprogramming, a far simpler procedure, could deliver comparable benefits, and how the depth of reprogramming affects subsequent recovery and effector function. HAECs were included because arterial endothelial dysfunction and senescence drive atherosclerosis and other age-associated cardiovascular disease^23,24^; we used adult arterial endothelium rather than umbilical vein endothelium, the source of much of the existing human endothelial literature, because adult donor-derived cells more directly represent the age context examined in this study.

After SeV-OSKM delivery, HDFs and CTLs underwent pronounced morphological changes and HAECs more modest ones, but all three cell types showed changes consistent with the initiation of reprogramming (Figure 1B). The kinetics of double-positive cells were cell-type-dependent—HDFs became GFP+OFP+ at earlier time points than the other cell types—but by days 5-7 after infection, roughly 30%-60% of cells were double-positive across the three cell types (Figure S1A). To analyze partial reprogramming in greater detail, we first monitored markers representative of somatic identity and early reprogramming for each cell type: CD13 and SSEA5 for HDFs, CD8α and SSEA4 for CTLs, and CD31 and SSEA4 for HAECs. After gating on GFP+OFP+ cells by flow cytometry, we defined three groups for each cell type according to the expression levels of these two markers (group 1 (Gr1): high-low, group 2 (Gr2): low-low, and group 3 (Gr3): low-high) (Figure 1C). We then examined the temporal changes in the proportions of these groups after SeV-OSKM transduction (Figure 1D). HDFs and CTLs both transitioned from Gr1 to Gr2 and subsequently accumulated in Gr3. CTLs progressed furthest over the measured interval: Gr2 exceeded Gr1 by day 5, and Gr3 became the predominant population by day 9. In contrast, HAECs progressed rapidly from Gr1 to Gr2, but rarely to Gr3 even after more than 10 days of reprogramming. Consistent with these progression patterns, iPSC derivation efficiency differed markedly by cell type: 3% for CTLs and 1% for HDFs; HAECs could generate iPSCs from all donors (Figure S1C) but with a very low mean efficiency of approximately 0.02% (Figure 1E).

Next we examined the transcriptomes and DNA methylation profiles of the partially reprogrammed cells. We chose time points at which we could recover a relatively balanced sampling of cells from each group: day 7 for HDFs and HAECs and day 5 for CTLs, because later time points became increasingly dominated by advanced states. Each sorted HDF and HAEC population was replated separately. CTLs were maintained in adherent culture, as in the iPSC derivation protocol, from the day after SeV infection until sorting, after which each sorted CTL population was immediately transferred to a separate suspension culture. At the same time, the incubation temperature was shifted from 35°C to 38°C to suppress replication of the temperature-sensitive SeV genome and thereby terminate SeV-OSKM expression. In parallel, nucleic acids were extracted from each sorted group for bulk RNA-seq and DNA methylation array profiling (Figure 1F). Because HAEC Gr3 cells were extremely rare, continued culture and subsequent sampling were not feasible, and HAEC Gr3 was excluded from downstream analyses.

Principal component analysis (PCA) of bulk RNA-seq showed clear separation of samples by group in all cell types and donors, confirming that the marker-defined groups represent substantially different reprogramming states (Figure S1D). To compare two key early events in reprogramming—loss of somatic identity and acquisition of an epithelial-associated program, a hallmark of the initiation phase of fibroblast reprogramming^25,26^—we calculated a cell type-specific identity module score from representative somatic markers and an epithelial-associated module score derived from MET-associated genes (Figure 1G). In HDFs, the epithelial-associated module captures the canonical MET program of fibroblast reprogramming; in CTLs and HAECs, it denotes acquisition of the same epithelial-associated gene set but does not imply a literal mesenchymal-to-epithelial transition. In HDFs, identity module scores decreased markedly from Gr1 to Gr2, whereas epithelial-associated module scores, which were already high in Gr1 relative to Ctrl HDFs, rose progressively through Gr2 and Gr3. CTLs also showed a marked reduction in identity module scores from Gr2 onward, although the additional decrease from Gr2 to Gr3 was limited. Epithelial-associated module scores were overall lower in CTLs than in HDFs but increased from Gr1 to Gr3. HAECs likewise showed a decrease in the identity module score and an increase in the epithelial-associated module score from Gr1 to Gr2.

We next performed DNA methylation array profiling and estimated DNA methylation age using epigenetic clocks (Figure 1H). For HDFs and HAECs, we used the Horvath clock^4^. Both HDFs and HAECs showed a reduction in DNA methylation age from Gr1 to Gr2, and in HDFs from Gr2 to Gr3. In HAECs, DNA methylation age was already strongly reduced in Gr1, nearly to the level observed in Gr2. In CTLs, the Horvath clock produced large variability between samples and did not clearly capture OSKM-induced reprogramming progression (Figure S1E). We therefore also evaluated the BLUP clock (Figure 1H)^27^ and IntrinClock (Figure S1F)^28^, which have been reported for DNA methylation age estimation in blood-lineage samples. Among these, the BLUP clock showed relatively limited sample-to-sample variation in our CTL samples and captured a reduction in predicted age consistent with transcriptomic progression. The reduction in DNA methylation age in CTL Gr1 and Gr2 was limited compared with the reductions observed in HDFs and HAECs, but Gr3 showed a marked decrease. In summary, we observed lineage-specific rates of somatic-identity loss, epithelial-associated reprogramming, pluripotency acquisition, and DNA methylation age reduction, indicating that OSKM triggers different reprogramming kinetics in different lineages.

### Reporter clearance and recovery kinetics vary by cell type and reprogramming depth

Next we aimed to analyze the kinetics and extent of recovery after OSKM withdrawal. Immediately after sorting, cultures were shifted from 35°C to 38°C to inhibit replication of the temperature-sensitive SeV genomes and initiate the loss of vector-derived OSKM expression. We defined reporter clearance as the point at which GFP and OFP fluorescence returned to the level of non-transduced cells, measured below (Figures 2A–2C). One day after the shift, while both reporters were still readily detectable, HDF Gr1 cells retained a spindle-like morphology similar to Ctrl cells, whereas HDF Gr2 and Gr3 cells were more rounded. HAEC Gr2 cells were also more rounded than Gr1 cells, although the difference was less pronounced. CTL morphology varied little among groups, but fewer Gr3 cells remained (Figure S2A). We conclude that morphological recovery had barely begun after one day at 38°C.

**Figure 2.**
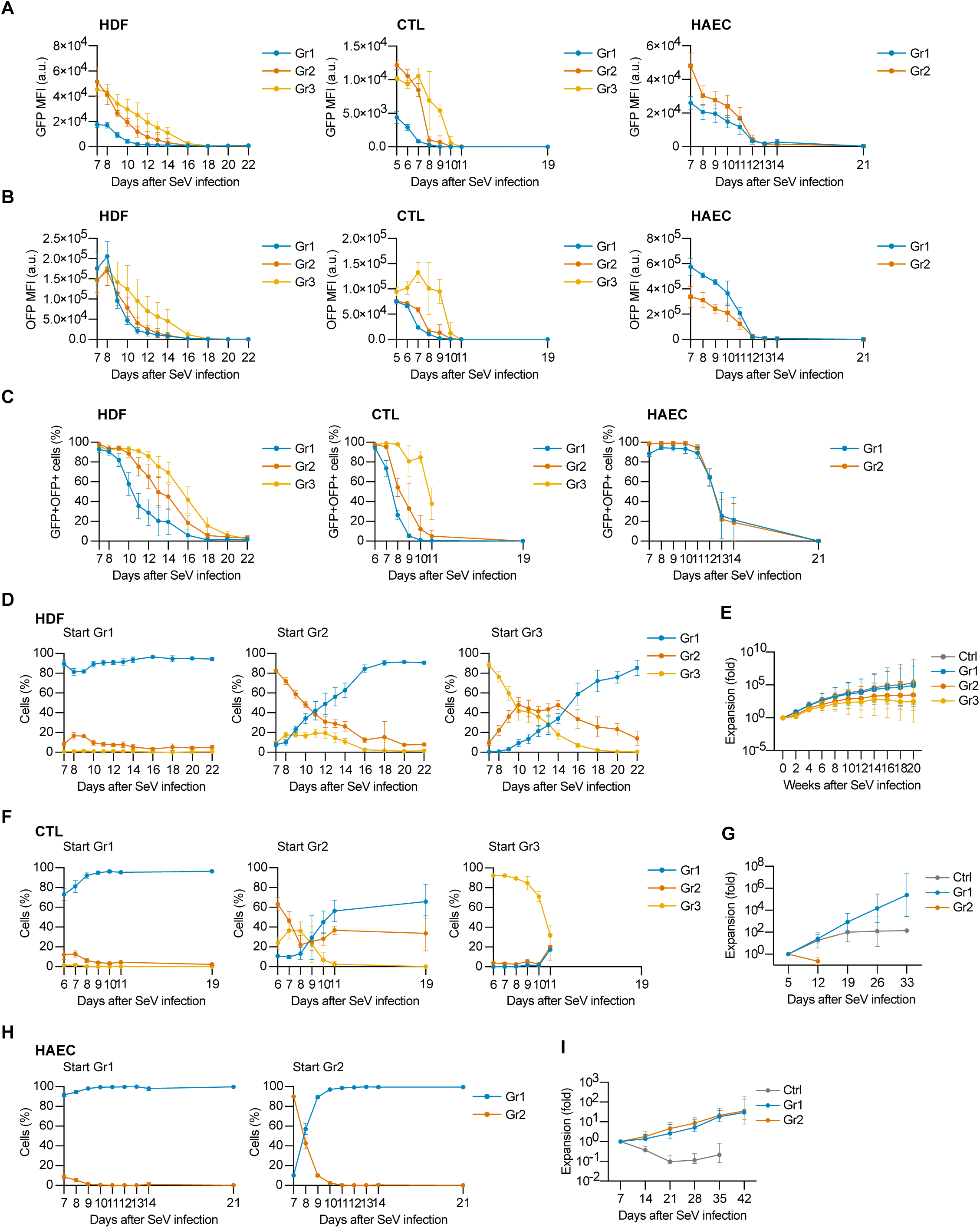
Reprogramming factor withdrawal reveals cell-type-and state-dependent reversibility. (A and B) Time-course analysis of GFP and OFP median fluorescence intensity (MFI) after temperature shift in sorted HDF, CTL, and HAEC groups. GFP and OFP report SeV-KOS-GFP and SeV-cMYC-OFP abundance, respectively. Line plots show mean values from four independent donors with SD. The groups shown are indicated in each panel: HDF and CTL panels show Gr1, Gr2 and Gr3, whereas HAEC panels show Gr1 and Gr2 only, because HAEC Gr3 was too rare to culture (Figure 1D). In (A)– (C), the CTL Gr3 traces end early because those cells were lost soon after sorting. (C) Time-course quantification of GFP+OFP+ cells after temperature shift, gated relative to non-SeV-transduced control cells. Line plots show mean values from four independent donors with SD. (D) Marker-defined state transitions after sorting of HDF Gr1, Gr2, and Gr3 cells. The plots show how cells sorted into each group redistributed among Gr1, Gr2, and Gr3 after withdrawal of reprogramming factors. Data are from four independent donors and are shown as mean ± SD. (E) Long-term expansion of sorted HDF populations after temperature shift. Data are shown as fold expansion over time, with mean ± SD from four independent donors. (F) Marker-defined state transitions after sorting of CTL Gr1, Gr2, and Gr3 cells. CTL Gr3 cells were lost early because of poor proliferative recovery. Data are mean ± SD from four independent donors. (G) Expansion of CTL groups after return to suspension culture. Cells were restimulated with CD3/CD28 beads weekly at the indicated time points. Gr1 maintained robust expansion, whereas Gr2 failed to sustain growth; Gr3 was not included because it was lost earlier. Data are mean ± SD from four independent donors. (H) Marker-defined state transitions after sorting of HAEC Gr1 and Gr2 cells. HAEC Gr2 rapidly converted to a Gr1-like state after temperature shift. Data are mean ± SD from four independent donors. (I) Expansion of HAEC Ctrl, Gr1, and Gr2 cells after temperature shift. Both Gr1 and Gr2 expanded more robustly than Ctrl after reprogramming factor withdrawal. Data are mean ± SD from four independent donors.

To monitor the loss of SeV-derived reporter expression after the temperature shift, we quantified GFP and OFP median fluorescence intensity (MFI) over time. At the time of sorting, GFP MFI, reflecting KOS expression, was higher in Gr2 and Gr3 populations than in Gr1 across all cell types (Figure 2A). After the temperature shift, HDF Gr1 cells showed the fastest decline in MFI and reached the baseline level of non-SeV-transduced cells by day 14 (7 days after the temperature shift), followed by Gr2 and Gr3. CTLs showed the most rapid reduction in MFI overall. CTL Gr1 declined fastest, and by day 9 (4 days after the temperature shift), both Gr1 and Gr2 reached MFI levels comparable to non-SeV-transduced cells. CTL Gr3 cells proliferated extremely poorly and disappeared before the MFI reached baseline. The more rapid fluorescence loss in CTLs than in HDFs or HAECs was likely due in part to stronger signal dilution through cell proliferation. In HAECs, Gr1 and Gr2 showed similar rates of MFI decline and reached the baseline slightly earlier than in HDFs, around day 12 (5 days after the temperature shift). In contrast to KOS-GFP MFI, OFP MFI, reflecting cMYC expression, tended to be higher in Gr1 than in Gr2 or Gr3, but the post-temperature-shift decline pattern was broadly similar to that of GFP (Figure 2B).

We also followed the fraction of double-positive (GFP+OFP+) cells (Figure 2C) and found that they disappeared fastest in Gr1, followed by Gr2 and Gr3, in both HDFs and CTLs. CTL Gr3 cells were lost before later measurements because of poor proliferation. HAEC Gr1 and Gr2 lost reporter signal with similar kinetics to each other; HAEC Gr3 was too rare to culture and was not analyzed. Thus, GFP and OFP signals declined gradually after temperature shift in every analyzable condition, albeit with cell-type-and state-dependent kinetics, consistent with progressive loss of vector-derived OSKM expression. We next related reporter-clearance kinetics to recovery of lineage-marker states and proliferative capacity. In HDFs, cells sorted as Gr1 remained almost entirely in a Gr1 state thereafter. By contrast, cells sorted as Gr2 or Gr3 gradually changed group profile: by day 22, nearly all Gr2 cells had returned to Gr1, and 10%–20% of Gr3 cells were classified as Gr2 because of relatively weak CD13 expression, indicating delayed recovery of HDF identity (Figure 2D and Figure S2B). We also followed HDF proliferation after the temperature shift over the long term (Figure 2E). Ctrl and Gr1 expanded at similar rates, whereas Gr2 and Gr3 proliferated more slowly than Ctrl during the first two weeks after temperature shift. Gr2 later showed partial recovery, but Gr3 remained clearly growth-impaired even 20 weeks after SeV infection. We did not observe overt signs of cell death, such as increased floating cells, in any group after temperature shift.

In CTLs, most Gr1 cells maintained their Gr1 state after temperature shift, similar to HDF Gr1. Cells sorted as Gr2 became predominantly Gr1 by day 19 but still contained approximately 30% Gr2-like cells. Gr3 cells exhibited extremely poor proliferation after the temperature shift, remained Gr3-dominant, and were lost after day 11 (Figure 2F and Figure S2C). Longitudinal analysis of cell expansion showed that Gr2 also proliferated poorly and failed to sustain expansion. In contrast, Gr1 expanded better than Ctrl even after four consecutive weekly rounds of CD3/CD28 stimulation following cell sorting (Figure 2G).

In HAECs, Gr2 cells reverted to a Gr1-like state far faster than the loss of GFP+OFP+ signal alone would predict; by day 10 (3 days after the temperature shift) nearly all Gr2 cells had converted to Gr1 (Figure 2H and Figure S2D), and both Gr1 and Gr2 sustained robust proliferation compared with Ctrl (Figure 2I).

Together, these experiments defined two features of the recovery phase—the time required for SeV-derived reporter signals to fall below detection and the kinetics with which lineage identity and proliferative capacity recover—and showed that both varied by cell type and reprogramming depth.

### Post-clearance outcomes reveal lineage-specific relationships between reprogramming benefits and functional recovery

Having established reporter-clearance and recovery kinetics, we next analyzed cells after GFP and OFP signals became undetectable. For each cell type, we asked whether the cells retained the reduction in DNA methylation age induced by reprogramming and whether they recovered their initial lineage identity and specialized function at the defined endpoint (Figure 3).

**Figure 3.**
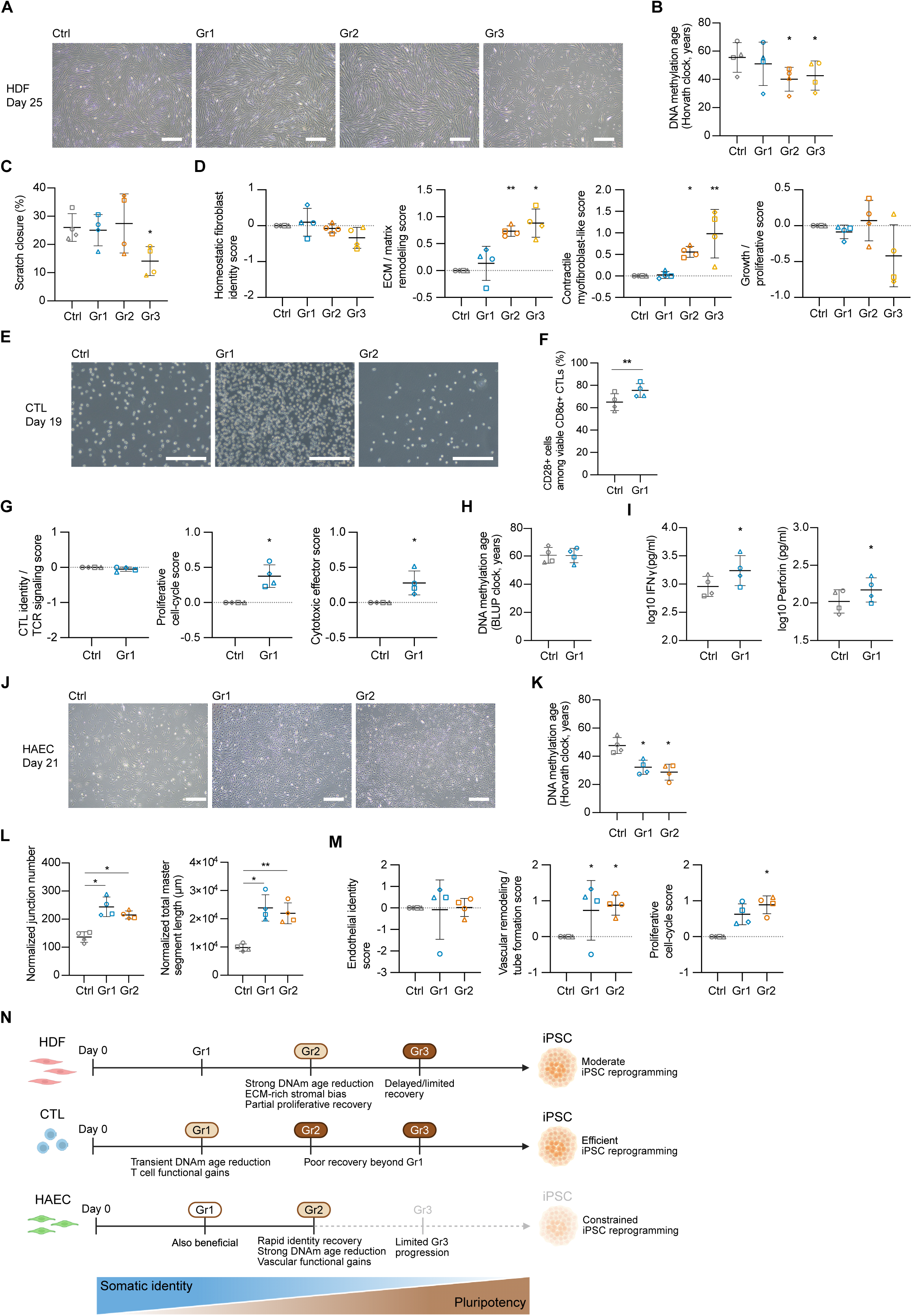
Distinct reprogramming trajectories produce cell type-specific recovery and functional outcomes. (A) Representative phase-contrast images of HDF Ctrl, Gr1, Gr2, and Gr3 at day 25 after SeV infection, following withdrawal of reprogramming factors and loss of fluorescent SeV signals. Scale bars, 200 µm. (B) DNA methylation age of HDFs at the endpoint, estimated using the Horvath clock. The central horizontal line indicates the mean of four independent donors, and error bars indicate SD; open symbols indicate HDF donors circle 54F, square 48F, triangle 56M, and diamond 48M. Repeated-measures one-way ANOVA followed by Dunnett’s multiple comparisons test versus Ctrl was used. \**p* < 0.05; unmarked comparisons are not significant. (C) Quantification of scratch wound closure in HDFs. The central horizontal line indicates the mean of four independent donors, and error bars indicate SD; open symbols indicate the HDF donors listed in panel B. Repeated-measures one-way ANOVA followed by Dunnett’s multiple comparisons test versus Ctrl was used. \**p* < 0.05; unmarked comparisons are not significant. (D) HDF endpoint bulk RNA-seq module scores for homeostatic fibroblast identity, ECM/matrix remodeling, a contractile myofibroblast-like state, and growth/proliferation. For visualization, donor-level module scores are shown as differences from the matched Ctrl score; statistical comparisons and displayed significance values were calculated from the corresponding original (uncentered) donor-level scores. The central horizontal line indicates the mean of four independent donors, and error bars indicate SD; open symbols indicate the HDF donors listed in panel B. Repeated-measures one-way ANOVA followed by Dunnett’s multiple comparisons test versus Ctrl was used for multi-condition comparisons. \**p* < 0.05, \*\**p* < 0.01, \*\*\**p* < 0.001; unmarked comparisons are not significant. (E) Representative phase-contrast images of CTL Ctrl, Gr1, and Gr2 at day 19 after SeV infection. Scale bars, 200 µm. (F) Flow cytometric quantification of CD28+ cells among viable CD8α+ CTLs at day 19. The central horizontal line indicates the mean of four independent donors, and error bars indicate SD; open symbols indicate CTL donors circle 58M, square 55M, triangle 56F, and diamond 54F. Ctrl and Gr1 were compared using a two-tailed paired t test. \*\**p* < 0.01. (G) Bulk RNA-seq module scores at day 19 for CTL identity/TCR signaling, proliferative cell-cycle state, and cytotoxic effector function. For visualization, donor-level module scores are shown as differences from the matched Ctrl score; the paired statistical comparisons and displayed significance values were calculated from the corresponding original (uncentered) donor-level scores. The central horizontal line indicates the mean of four independent donors, and error bars indicate SD; open symbols indicate the CTL donors listed in panel F. Ctrl and Gr1 values were compared using a two-tailed paired t test. \**p* < 0.05; unmarked comparisons are not significant. (H) DNA methylation age of CTLs at day 26, estimated using the BLUP clock. The central horizontal line indicates the mean of four independent donors, and error bars indicate SD. A two-tailed paired t test was used; unmarked comparisons are not significant. (I) Log10-transformed IFN-γ and perforin concentrations in CTL culture supernatants collected on day 30 after SeV-OSKM transduction, four days after CD3/CD28 restimulation on day 26. Open symbols indicate individual CTL donors, with donor identities indicated as in panel F. The central horizontal line indicates the mean, and error bars indicate SD. Ctrl and Gr1 values were compared using a two-tailed paired t test. \**p* < 0.05. (J) Representative phase-contrast images of HAEC Ctrl, Gr1, and Gr2 at day 21 after SeV infection. Scale bars, 200 µm. (K) DNA methylation age of HAECs at day 21, estimated using the Horvath clock. The central horizontal line indicates the mean of four independent donors, and error bars indicate SD; open symbols indicate HAEC donors circle 63M, square 61M, triangle 51F, and diamond 50F. Repeated-measures one-way ANOVA followed by Dunnett’s multiple comparisons test versus Ctrl was used. \**p* < 0.05. (L) Cell-number-normalized junction number and total master segment length in HAEC tube-formation assays. The remaining normalized metrics, segment number and master segment number, are shown in Figure S3E. The central horizontal line indicates the mean of four independent donors, and error bars indicate SD; open symbols indicate the HAEC donors listed in panel K. Repeated-measures one-way ANOVA followed by Dunnett’s multiple comparisons test versus Ctrl was used. A prespecified comparison between Gr1 and Gr2 was additionally performed using a Sidak-adjusted comparison; no significant difference was detected. \**p* < 0.05, \*\**p* < 0.01; unmarked comparisons are not significant. (M) HAEC endpoint bulk RNA-seq module scores for endothelial identity, vascular remodeling/tube formation, and proliferative cell-cycle state. For visualization, donor-level module scores are shown as differences from the matched Ctrl score; statistical comparisons and displayed significance values were calculated from the corresponding original (uncentered) donor-level scores. The central horizontal line indicates the mean of four independent donors, and error bars indicate SD; open symbols indicate the HAEC donors listed in panel K. Repeated-measures one-way ANOVA followed by Dunnett’s multiple comparisons test versus Ctrl was used. \**p* < 0.05; unmarked comparisons are not significant. (N) Summary model integrating marker-defined progression, recovery phenotypes, and competence for full iPSC reprogramming across the three cell types. Light brown highlighting marks states with comparatively favorable post-withdrawal outcomes under the tested conditions: CTL Gr1 and HAEC Gr1/Gr2 showed functional improvements, whereas HDF Gr2 showed a mixed but comparatively balanced outcome, retaining DNA methylation age reduction while avoiding the marked recovery impairment observed after Gr3. Darker brown marks states associated with impaired recovery in HDFs and CTLs, and gray indicates HAEC Gr3/iPSC progression that was rarely achieved and was not evaluated in post-sort assays. The model emphasizes that correspondingly ordered states have different biological consequences across cell types and that recovery outcomes do not scale directly with full iPSC reprogramming efficiency.

In HDFs, reduced DNA methylation age persisted despite incomplete functional recovery. By day 25, HDFs recovered a morphology broadly similar to Ctrl, and their global transcriptomes converged toward a shared somatic state (Figures 3A and S3B). Gr2 and Gr3 nevertheless retained significantly lower DNA methylation age estimates than Ctrl, whereas Gr1 was variable across donors and no longer differed significantly from Ctrl (Figure 3B). Homeostatic fibroblast identity was broadly reacquired, but Gr2 and Gr3 retained elevated ECM/matrix-remodeling and contractile myofibroblast-like programs (Figure 3D).

Functional recovery depended on reprogramming depth: scratch closure varied among Gr2 donors and did not differ significantly from Ctrl, whereas it was significantly reduced in Gr3, consistent with Gr3 cells’ persistent growth impairment (Figures 3C and S3A). Among the HDF states tested, Gr2 therefore provided the most balanced outcome: sustained DNA methylation age reduction without the severe growth and wound-closure deficits of Gr3. Nevertheless, proliferative recovery remained incomplete and stromal remodeling persisted, indicating a mixed outcome rather than complete functional normalization.

In CTLs, sustained recovery was restricted to the shallow Gr1 state because Gr2 and Gr3 cells failed to sustain expansion after OSKM suppression. At day 19, the remaining Gr1 cells resembled Ctrl morphologically, their global transcriptomes had converged toward Ctrl, and CTL identity scores did not differ significantly (Figures 3E, 3G, and S3C). Gr1 nevertheless contained a higher fraction of CD28+ cells than Ctrl CTLs did, and showed increased proliferative cell-cycle and cytotoxic-effector scores (Figures 3F and 3G). By day 26, the DNA methylation age reduction observed at sorting was no longer detectable by the BLUP, Horvath, or IntrinClock estimates (Figures 3H and S3D). After restimulation, however, Gr1 produced more IFN-γ and perforin than did Ctrl CTLs at day 30 (Figure 3I). Thus, shallow reprogramming of CTLs allowed robust recovery combined with enhanced effector-associated function, but not the maintenance of DNA methylation age reduction.

In HAECs, Gr1 and Gr2 rapidly recovered identity and showed improved vascular function. By day 21, both states showed no major morphological differences from Ctrl, had converged toward Ctrl in global transcriptomic space, and achieved comparable endothelial identity (Figures 3J, 3M, and S3G). Both states retained a low DNA methylation age, with a larger numerical reduction in Gr2 (Figure 3K). Gr1 and Gr2 also improved all four cell-number-normalized tube-formation metrics and increased vascular-remodeling/tube-formation programs, and the proliferative cell-cycle score increased significantly in Gr2 (Figures 3L, 3M, and S3E–S3F). Although Gr1 and Gr2 did not differ significantly, Gr2 showed the most consistent profile across donors, combining rapid identity recovery with sustained molecular-age reduction and markedly improved vascular function.

Together, these post-clearance endpoints revealed distinct relationships among reprogramming depth, persistence of the DNA methylation age reduction, and functional recovery among the 3 lineages (Figure 3N). Deeper HDF reprogramming states retained DNA methylation age reduction but showed incomplete functional recovery; CTLs were only recoverable after shallow progression and did not retain the initial age reduction; and HAEC Gr1 and Gr2 recovered identity while retaining molecular-age reduction and improving vascular function. Importantly, reprogramming states that were comparable between cell types (e.g., Gr2 or Gr1) did not produce equivalent relationships between epigenetic rejuvenation and functional recovery.

### The reprogramming trajectories of HDFs, CTLs and HAECs share a TOX-associated regulatory node

We reasoned that the cell type-specific responses to OSKM exposure and subsequent withdrawal reflected different molecular trajectories between cell lineages during the reprogramming process. To test this hypothesis, we performed single-nucleus multiome analysis on cells from the sorted groups on day 7 for HDFs and HAECs and day 5 for CTLs (Figure 4A). RNA-based integrated UMAP separated the three cell types and resolved Day 0, Ctrl and the sorted groups within each cell type (Gr1–Gr3 for HDFs and CTLs, Gr1 and Gr2 for HAECs) (Figure 4B). We then applied UniTVelo RNA velocity analysis^29^ to the sorted Gr populations, specifying Gr1 cells as the starting population to orient the inferred trajectories. Inferred velocity streamlines extended from Gr1 toward Gr2 in HAECs and toward the Gr2/Gr3 domains in HDFs and CTLs (Figure 4C, left). UniTVelo latent time showed concordant gradients, with lower values concentrated in Gr1 and higher values extending into the deeper groups (Figure 4C, upper right). These findings confirm that in each lineage, the analyzed groups represent a continuous progression from shallow to more advanced reprogramming states.

**Figure 4.**
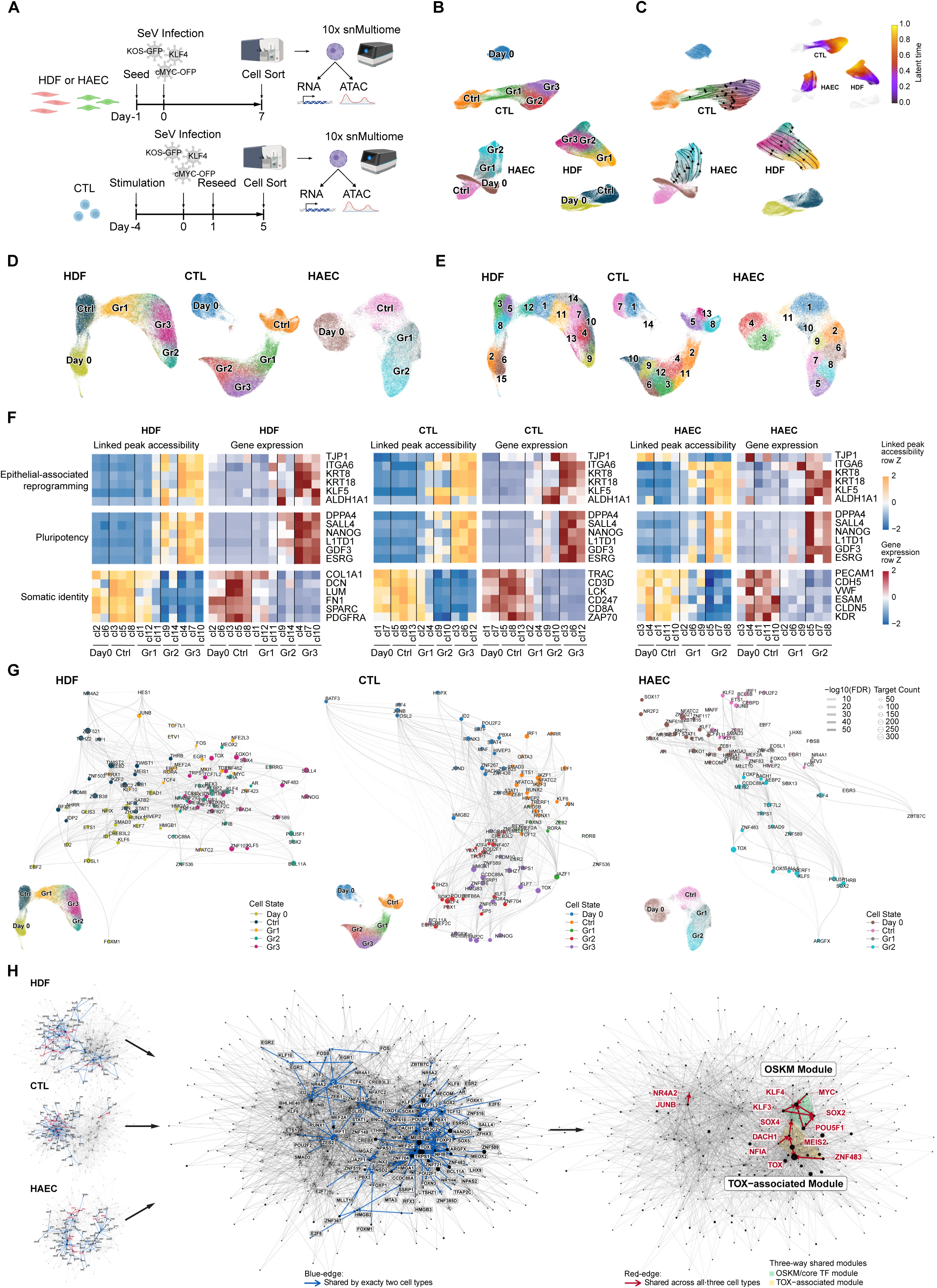
Divergent lineage trajectories converge on a shared Pando-inferred TOX-associated regulatory node. (A) Experimental overview for single-nucleus multiome profiling. HDFs and HAECs were profiled on day 7 and CTLs on day 5 after SeV-OSKM infection. (B) RNA-based integrated UMAP of HDF, CTL, and HAEC cells across Day 0, Ctrl, Gr1, Gr2, and Gr3 states. Cells segregated primarily by cell type, with additional separation among experimental states within each cell type. (C) UniTVelo RNA-velocity and latent-time analyses of marker-defined reprogramming groups projected onto RNA-based UMAPs. Left, inferred velocity streamlines, with Gr1 cells specified as roots, extend toward Gr2 in HAECs and toward the Gr2/Gr3 domains in HDFs and CTLs. Upper right, latent-time projections show concordant within-cell-type gradients from lower values in Gr1 toward higher values in more advanced states. Gray points indicate Day 0 and Ctrl cells shown for reference but not included in the cell type-specific UniTVelo fitting. Latent time was inferred and scaled separately within each cell type and is therefore not quantitatively comparable across cell types. (D) Cell type-specific ATAC-based UMAPs showing chromatin-state organization of Day 0, Ctrl, and reprogramming groups. (E) Leiden clustering used to define representative clusters for downstream visualization of linked peak accessibility, gene expression, and gene regulatory networks. Representative clusters were selected from the most abundant clusters within each group. (F) Linked peak accessibility and gene expression heatmaps for representative epithelial-associated reprogramming (MET-associated), pluripotency, and somatic identity genes in HDFs, CTLs, and HAECs. Values are row-scaled Z-scores for linked peak accessibility and gene expression, respectively. (G) Cell type-specific Pando regulatory networks. Node color indicates the dominant cell state assigned to each transcription factor, node size indicates the number of inferred target genes, and edge darkness represents −log10(FDR), with darker edges indicating greater statistical significance. TOX-associated regulatory nodes were identified in more advanced reprogramming states across all three cell types. (H) Shared Pando regulatory networks across cell types. Blue edges indicate regulatory edges shared by exactly two cell types, whereas red edges indicate regulatory edges shared across HDFs, CTLs, and HAECs. The three-way intersection recovered the canonical OSKM/core TF module and a distinct TOX-associated module. All displayed edges are Pando-inferred regulatory relationships and do not establish direct TF occupancy or causal regulation.

For more detailed multiome analysis, we generated ATAC-based UMAPs for each cell type. In each case, groups separated clearly along reprogramming progression (Figure 4D). Because grouping was defined by flow cytometry gating, especially for Gr1, Gr2, and Gr3, minor populations with features spanning more than one group can occur. To minimize the impact of such cells and clarify group-level features, we performed clustering such that each group was divided into roughly one to four clusters (Figure 4E), and we selected the two to three representative clusters with the largest cell fractions in each group for downstream state-resolved visualization and module summaries (Figure S4A).

Linked peak accessibility and gene expression showed an increasing loss of somatic identity and gains in epithelial-associated reprogramming and pluripotency programs, with the strongest changes in Gr2 and Gr3 in HDFs and CTLs and in Gr2 in HAECs (Figure 4F). Pluripotency-gene expression was most pronounced in CTL Gr3. In contrast, HAEC Gr2 displayed relatively low pluripotency-gene expression compared with its epithelial-associated reprogramming program and with the more progressed states of the other cell types (Figure S4B). HDFs and HAECs also retained somatic identity-associated expression more strongly than CTLs. These patterns paralleled the measured iPSC derivation efficiencies and were consistent with a bottleneck in HAEC progression after the early response and epithelial-associated activation, between Gr2 and the Gr3/iPSC states. Thus, progression toward pluripotency retained strong lineage-specific features.

This observation, together with the distinct recovery and functional phenotypes of the different lineages upon OSKM removal, raised the question of whether the three trajectories relied on entirely separate regulatory machinery. To answer this question, we performed Pando gene regulatory network (GRN) analysis^30^ separately in HDFs, CTLs, and HAECs and compared the inferred networks across lineages. We first generated state-resolved views of each cell type-specific network in relation to the corresponding ATAC UMAP. In both this analysis and the subsequent cross-lineage comparison, the displayed networks were restricted to transcription factors: node size represented the number of inferred targets, whereas the number of edges connected to a node reflected the number of inferred regulatory interactions. Somatic-identity-and pluripotency-associated TF networks were distributed according to reprogramming progression, with pluripotency-related networks becoming increasingly prominent in progressed states (Figure 4G). These patterns were consistent with the flow-cytometric and bulk RNA-seq evidence of progressive reprogramming, but the complexity of the individual cell type-specific networks did not by itself nominate a common regulator.

We therefore intersected inferred TF regulatory edges across cell types to identify conserved network structure. Edges shared by exactly two cell types formed an extensive and highly interconnected network in which discrete regulatory cores could not be readily prioritized. By contrast, restricting the analysis to edges shared across all three cell types sharply reduced network complexity and revealed a compact cross-lineage core. This core contained a canonical reprogramming-factor module comprising all four OSKM factors—OCT4 (*POU5F1*), SOX2, KLF4, and MYC—as well as a smaller JUNB-associated component and a distinct module organized around TOX (Figure 4H and Figure S4C). Within the network shared across all three cell types, TOX was the largest node, reflecting the highest inferred target count among the displayed TFs, and was linked to multiple TFs by regulatory edges shared across all three cell types. We therefore termed this network the Pando-inferred TOX-associated module. Recovery of the established OSKM module provided an internal positive control for the biological relevance of the three-way network intersection, whereas the prominence of TOX nominated it as an unexpected candidate regulator shared across lineages during reprogramming. TOX (thymocyte selection-associated HMG box) is a nuclear HMG-box protein of a subfamily thought to bind DNA largely through structure rather than sequence, and is best characterized in T-cell development and exhaustion; its paralog *TOX4* has been implicated in early murine reprogramming, but TOX itself has no described role in human somatic cell reprogramming^31,32^. We therefore prioritized it for functional analysis.

### *TOX* is transiently induced across the three cell types within a candidate OSK-responsive cis-regulatory landscape

We first examined *TOX* expression during reprogramming. Low basal *TOX* expression was detectable in both Day 0 and Ctrl populations of HDFs and HAECs, and increased markedly in later reprogramming states, particularly from Gr2 onward. In CTLs, *TOX* induction was most prominent in Gr3, although *TOX*-high cells (cl6 and cl12) were also substantially represented in Gr2 (Figure S4A). Consistent with these expression patterns, accessibility of *TOX*-linked peaks and *TOX* RNA expression were highest in HDF Gr2/Gr3, CTL Gr3, and HAEC Gr2 (Figures 5A and 5B).

**Figure 5.**
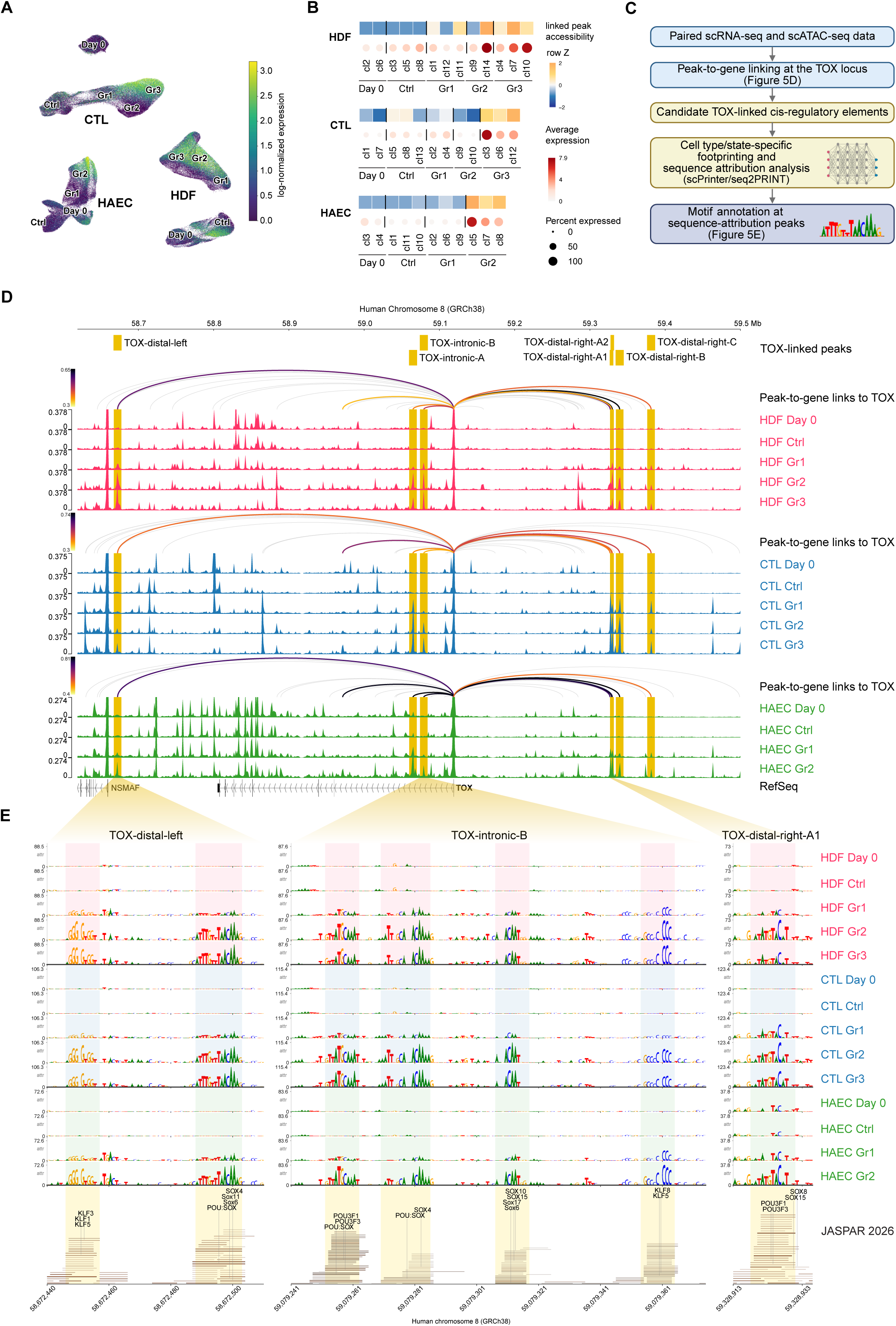
Transient *TOX* induction across three cell types accompanies opening of candidate OSK-responsive cis-regulatory elements. (A) *TOX* RNA expression projected onto single-nucleus multiome UMAPs across HDF, CTL, and HAEC reprogramming states. (B) Accessibility of peaks linked to *TOX* and *TOX* RNA expression across selected clusters representing each reprogramming state. Linked-peak accessibility is shown as row-scaled Z-scores. For *TOX* expression, dot color indicates average log-normalized expression and dot size indicates the percentage of *TOX*-expressing cells. (C) Analysis workflow used to interrogate upstream regulation of *TOX*. Peak-to-gene links were used to nominate *TOX*-linked CREs, followed by cell type-and state-resolved scPrinter/seq2PRINT footprint and sequence-attribution analysis and motif annotation at sequence-attribution peaks. The workflow explicitly links the locus-level peak-to-gene analysis to panel D and the motif-annotated sequence-attribution analysis to panel E. (D) Enlarged genome browser view of the *TOX* locus showing chromatin accessibility tracks, peak-to-gene links to *TOX*, and seven labeled intronic and distal *TOX*-linked candidate CREs in HDFs, CTLs, and HAECs. Tracks are shown for Day 0, Ctrl, and the indicated reprogramming groups. (E) Motif-centered scPrinter/seq2PRINT sequence-attribution profiles at three selected *TOX*-linked candidate CREs across HDFs, CTLs, and HAECs. Corresponding JASPAR 2026 motif matches are shown below each interval; the four remaining labeled candidate CREs are shown in Figure S5B. Reprogramming-state-dependent attribution peaks contained numerous POU-family, SOX-family, POU:SOX composite, and KLF-family motif matches, nominating candidate POU/SOX/KLF inputs to *TOX* rather than establishing factor-specific binding at these elements.

To determine whether *TOX* induction also occurs during full reprogramming to iPSCs, we reanalyzed three independent human HDF-to-iPSC single-cell RNA-seq datasets generated using SeV-OSKM, lentiviral OSKM, or mRNA OSKMLN (OSKM plus LIN28 and NANOG)^33–35^. *TOX* was transiently induced during early-to-mid reprogramming in all three datasets and subsequently declined toward the established iPSC state (Figure S5A). Thus, transient *TOX* induction is reproducible across distinct reprogramming platforms and is concentrated in the early-to-mid phase of reprogramming.

To investigate the cis-regulatory basis of *TOX* induction during partial reprogramming, we integrated peak-to-gene linkage at the *TOX* locus with nomination of candidate *TOX*-linked cis-regulatory elements (CREs), cell type-and state-resolved scPrinter/seq2PRINT^36^ footprint and sequence-attribution analysis, and motif annotation at attribution peaks (Figure 5C). Genome browser tracks identified seven labeled intronic and distal candidate CREs that gained accessibility with reprogramming progression and showed peak-to-gene links to *TOX* across HDFs, CTLs, and HAECs (Figure 5D).

Analysis of all seven elements—three representative CREs in the main figure and the remaining four in the supplement—revealed cell type- and state-dependent scPrinter/seq2PRINT sequence-attribution profiles, supporting coordinated opening of multiple intronic and distal *TOX*-linked CREs rather than regulation through a single proximal element. JASPAR 2026 annotation^37^ revealed numerous POU-family, SOX-family, POU:SOX composite, and KLF-family motif matches at attribution peaks. These findings are consistent with potential inputs from the POU/SOX/KLF network, including plausible contributions from the exogenous OSK factors (Figure 5E and Figure S5B).

Together, these observations establish transient *TOX* induction as a shared feature of reprogramming across all three cell types and show that it is accompanied by opening of multiple *TOX*-linked candidate CREs. POU-, SOX-, and KLF-family motif matches nominate potential inputs from the OSK network but do not establish factor-specific occupancy or direct activation. We therefore asked whether TOX was merely a marker of advanced reprogramming states or a functional component of progression.

### *TOX* depletion impairs reprogramming progression across the three cell types

To test whether TOX contributes functionally to reprogramming rather than merely marking advanced states, we suppressed *TOX* during reprogramming using siRNAs in HDFs and HAECs and shRNAs in CTLs. HDFs and HAECs were treated separately with each of three *TOX*-targeting siRNAs (siTOX #1– #3), with transfection repeated every 3 days. CTLs were transduced separately with each of three puromycin-selectable lentiviral vectors expressing distinct *TOX*-targeting shRNAs (shTOX #1–#3).

Bulk RNA-seq and parallel flow cytometric analyses of the marker-defined Gr states were performed on day 7 for HDFs and HAECs and day 5 for CTLs, and iPSC colony formation was assessed on day 14. A non-targeting siRNA (siCtrl) for HDFs and HAECs and a scrambled-sequence shRNA (shCtrl) for CTLs served as matched controls (Figure 6A). *TOX* knockdown efficiency was assessed on day 7 in HDFs and HAECs after siRNA transfections on days 1 and 4, and on day 5 after SeV-OSKM transduction in CTLs that had undergone lentiviral shRNA transduction and selection before reprogramming (Figure 6B). Knockdown efficiency differed among reagents. The same siRNA sequences were used in HDFs and HAECs, with siTOX #1 and #2 producing particularly efficient knockdown. In CTLs, shRNA-mediated knockdown was weaker overall, with shTOX #2 producing the greatest reduction. *TOX* knockdown sharply reduced iPSC derivation efficiency in all three cell types. In HDFs and HAECs, siTOX #1 and #2 nearly abolished colony formation; in CTLs, shTOX #2 produced the largest decrease (Figure 6C). qPCR on day 2 showed that *TOX* knockdown did not decrease total SeV RNA (Figure 6D), SeV-KOS RNA (Figure S6A), or SeV-cMYC RNA (Figure S6B). Lower early abundance of the exogenous reprogramming-factor transcripts is therefore unlikely to account for the reduced colony formation.

**Figure 6.**
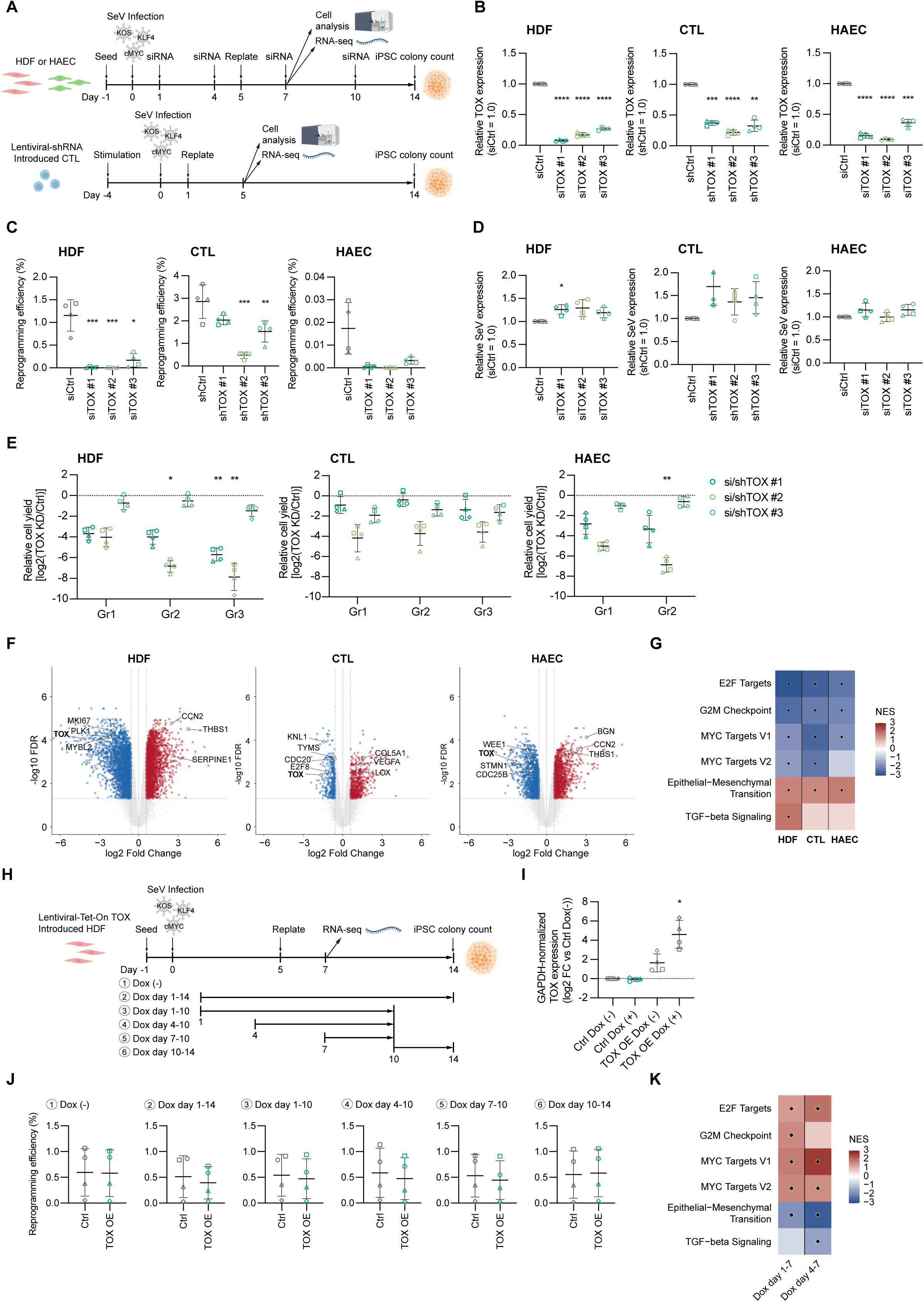
TOX supports reprogramming progression across three cell types. (A) Experimental design for *TOX* knockdown in HDFs, CTLs, and HAECs. HDFs and HAECs were treated with siRNAs during reprogramming, whereas CTLs were transduced with lentiviral shRNAs. Flow cytometric cell analysis and RNA-seq were performed on day 7 for HDFs and HAECs and day 5 for CTLs, and iPSC colonies were counted on day 14. (B) qPCR analysis of *TOX* knockdown efficiency on day 7 after SeV-OSKM transduction in HDFs and HAECs, following siRNA transfections on days 1 and 4, and on day 5 after SeV-OSKM transduction in CTLs transduced with lentiviral shRNAs before reprogramming. The central horizontal line indicates the mean of four independent donors, and error bars indicate SD; open symbols indicate individual donors: HDFs, circle 54F, square 48F, triangle 56M, diamond 48M; CTLs, circle 58M, square 55M, triangle 56F, diamond 54F; HAECs, circle 63M, square 61M, triangle 51F, diamond 50F. Repeated-measures one-way ANOVA followed by Dunnett’s multiple comparisons test versus siCtrl for HDFs and HAECs or shCtrl for CTLs was used for multi-condition comparisons. \*\**p* < 0.01, \*\*\**p* < 0.001, \*\*\*\**p* < 0.0001; unmarked comparisons are not significant. (C) iPSC reprogramming efficiency after *TOX* knockdown. The central horizontal line indicates the mean of four independent donors, and error bars indicate SD; open symbols indicate the donors listed in panel B. Repeated-measures one-way ANOVA followed by Dunnett’s multiple comparisons test versus the matched control (siCtrl for HDFs and HAECs; shCtrl for CTLs) was used. \**p* < 0.05, \*\**p* < 0.01, \*\*\**p* < 0.001; unmarked comparisons are not significant. (D) Total SeV RNA abundance after *TOX* knockdown, measured by qPCR on day 2 after SeV-OSKM infection. The absence of a decrease argues against lower total SeV RNA abundance at this time point as the explanation for reduced reprogramming efficiency. The central horizontal line indicates the mean of four independent donors, and error bars indicate SD; open symbols indicate the donors listed in panel B. Repeated-measures one-way ANOVA followed by Dunnett’s multiple-comparisons test versus the matched control (siCtrl for HDFs and HAECs; shCtrl for CTLs) was used where indicated. \**p* < 0.05; unmarked comparisons are not significant. (E) Relative yield of marker-defined Gr populations after *TOX* knockdown at day 7 for HDFs and HAECs and day 5 for CTLs. For each donor and Gr, yield is shown as log2(*TOX* KD/matched control), where the matched control was siCtrl for HDFs and HAECs and shCtrl for CTLs; absolute Gr-cell yields are shown in Figure S6E. The central horizontal line indicates the mean of four independent donors, and error bars indicate SD; open symbols indicate the donors listed in panel B. Relative Gr-cell yields were analyzed by two-way repeated-measures ANOVA with donor matching across Gr state and siRNA/shRNA identity and Geisser–Greenhouse correction. Post hoc comparisons among Gr states were performed separately for each *TOX*-targeting siRNA or shRNA. For HDFs and CTLs, the relative yields of Gr2 and Gr3 were compared with that of Gr1 using Dunnett-adjusted comparisons; for HAECs, the relative yield of Gr2 was compared with that of Gr1 using a Sidak-adjusted comparison. Asterisks above Gr2 or Gr3 denote significant differences from Gr1 for the same siRNA or shRNA and do not indicate that log2(KD/matched control) differs significantly from zero. \**p* < 0.05, \*\**p* < 0.01; unmarked comparisons are not significant. (F) Volcano plots of bulk RNA-seq after *TOX* knockdown in HDFs, CTLs, and HAECs. Blue and red points indicate significantly downregulated and upregulated genes, respectively, defined as FDR < 0.05 and |log2 fold change| > 1. Dashed lines indicate these thresholds. *TOX* and representative proliferation/cell-cycle genes are labeled among downregulated genes, whereas remodeling-or stress-associated genes are labeled among upregulated genes. (G) GSEA heatmap showing normalized enrichment scores (NES) for selected hallmark gene sets after *TOX* knockdown. E2F targets, G2M checkpoint, and MYC targets show negative enrichment, whereas EMT and TGF-β signaling show positive enrichment. Black dots indicate significantly enriched gene sets with FDR < 0.05. (H) Experimental design for doxycycline-inducible *TOX* overexpression in HDFs during reprogramming. HDFs carrying the lentiviral Tet-On *TOX* construct were induced with doxycycline at the indicated time windows, RNA was collected on day 7 for RNA-seq, and iPSC colonies were counted on day 14. (I) *TOX* expression after doxycycline induction. The central horizontal line indicates the mean of four independent donors, and error bars indicate SD; open symbols indicate HDF donors circle 54F, square 48F, triangle 56M, and diamond 48M. Repeated-measures one-way ANOVA followed by Dunnett’s multiple-comparisons test versus Ctrl Dox (+) was used where indicated. \**p* < 0.05; unmarked comparisons are not significant. (J) Reprogramming efficiency after *TOX* overexpression at different doxycycline induction windows. The central horizontal line indicates the mean of four independent donors, and error bars indicate SD; open symbols indicate the HDF donors listed in panel I. Ctrl and *TOX* OE values were compared using a two-tailed paired t test. No significant differences were detected. (K) GSEA heatmap after *TOX* overexpression. E2F target, G2M checkpoint, and MYC target gene sets show positive enrichment, whereas EMT and TGF-β signaling show negative enrichment, opposite to the pathway-level changes observed after *TOX* knockdown. Black dots indicate significantly enriched gene sets with FDR < 0.05. The corresponding individual-gene volcano plots are shown in Figure S6G.

To determine whether reduced iPSC derivation could reflect a toxic effect of TOX reduction, we reduced *TOX* expression in HDFs, HAECs, and CTLs, in the absence of SeV-OSKM. In HDFs and HAECs, *TOX* expression was measured 2 days after siRNA transfection and cell yields assessed on day 3. *TOX* expression was reduced by siTOX #1 and #2 in HDFs and by all three siRNAs in HAECs (Figure S6C). For CTLs, cells transduced with shTOX or shCtrl lentiviral vectors were selected with puromycin for 7 days, stimulated with CD3/CD28, and analyzed 3 days later. *TOX* expression remained very low in CTLs under this pre-reprogramming stimulation condition, resulting in variable relative estimates; however, the same shRNAs suppressed OSKM-induced *TOX* expression (Figure 6B). Cell yield was not significantly reduced in HDFs or CTLs. In HAECs, it was unaffected by siTOX #1 or #3 but was reduced by siTOX #2 (Figure S6D). Because siTOX #1 and #3 reduced *TOX* expression in HAECs without reducing cell yield, this effect of siRNA#2 is unlikely to reflect a requirement for TOX in HAEC survival; it may instead arise from a sequence-specific off-target toxicity of siRNA#2, although a deeper *TOX* depletion beyond the cells’ viability threshold cannot be excluded.

Together, these findings indicated that the negative impact of TOX loss of function on iPSC derivation could not simply be explained by a reduction in cell yield. We therefore examined Gr-cell yields during reprogramming, as altered accumulation of these intermediates could contribute to the cell type-specific outcomes of partial reprogramming.

At our chosen analysis time point (day 7 for HDFs and HAECs; day 5 for CTLs), reducing TOX lowered the mean absolute yield for each Gr relative to matched controls and across all three cell types, although the magnitude and statistical significance varied among reagents and populations (Figure S6E). We then analyzed log2(KD/matched-control) ratios to determine whether the magnitude of this reduction differed among Gr states (Figure 6E). In HDFs, the deepest Gr3 state was disproportionately reduced by the more effective siRNAs, whereas the sensitivity of Gr2 varied among siRNAs. In CTLs, neither the Gr2 nor the Gr3 log2 ratio differed significantly from that of Gr1 within an individual shRNA after correction. In HAECs, both Gr1 and Gr2 were reduced, but preferential loss of Gr2 was observed only with siTOX #2 and was not reproduced by the other siRNAs. Thus, TOX supports the establishment or maintenance of graded reprogramming intermediates across all three cell types, but each state’s dependence on TOX is cell type- and reagent-dependent.

### *TOX* knockdown shifts shared proliferative and EMT/TGF-β programs but produces cell type-specific gene-level responses

In cells still expressing SeV-OSKM, bulk RNA-seq after acute *TOX* knockdown revealed the largest number of differentially expressed genes (DEGs) in HDFs, followed by HAECs and CTLs. Although the individual genes affected differed among cell types, the volcano plots showed a convergent attenuation of cell-cycle-associated programs. Representative downregulated genes included *MKI67*, *PLK1*, and *MYBL2* in HDFs; *KNL1*, *TYMS*, *CDC20*, and *E2F8* in CTLs; and *WEE1*, *STMN1*, and *CDC25B* in HAECs, collectively implicating impaired DNA synthesis, mitotic progression, checkpoint regulation, and microtubule dynamics following *TOX* depletion (Figure 6F). Consistent with these gene-level changes, gene set enrichment analysis showed negative enrichment of E2F targets, G2M checkpoint, and MYC target gene sets across all three cell types (Figure 6G).

Conversely, the Hallmark epithelial–mesenchymal transition (EMT) gene set was positively enriched in all three cell types, and TGF-β signaling was also positively enriched, most prominently in HDFs (Figure 6G). These pathway-level changes were accompanied by the induction of cell-type-dependent extracellular-matrix and matricellular genes, including *CCN2*, *THBS1*, and *SERPINE1* in HDFs; *COL5A1* and *LOX*, together with *VEGFA*, in CTLs; and *BGN*, *CCN2*, and *THBS1* in HAECs (Figure 6F). The transcriptional response to *TOX* knockdown was broadly reciprocal to the gene-set dynamics observed across representative single-nucleus multiome clusters spanning the graded reprogramming states, in which more advanced reprogramming populations generally exhibited increased proliferative programs and reduced EMT- and TGF-β-associated activity, although the timing and magnitude of these changes varied among cell types (Figure S6F).

Together, these loss-of-function data show that TOX is necessary for efficient reprogramming progression across all three cell types. Because reprogramming to pluripotency depends on cell division, and a high proliferation rate is a known requirement for efficient iPSC derivation^38,39^, the attenuation of proliferative programs after *TOX* depletion could in principle account for the reduced iPSC output on its own. Two observations indicate that the yield deficit occurs mainly in the reprogramming context. First, *TOX* knockdown in the absence of SeV-OSKM largely preserved cell yield (Figures S6C and S6D). Second, Gr-cell yields were reduced at every depth during reprogramming (Figure S6E). The proliferative deficit is therefore largely confined to cells undergoing OSKM-driven progression.

Two further observations argue that the colony deficit is not a simple consequence of reduced cell number. First, in HDFs, the two measures were not proportional across reagents: si*TOX* #3 reduced Gr-cell yields by less than three-fold at every depth yet reduced colony output roughly eight-fold, whereas si*TOX* #2 reduced Gr3 yield by more than 100-fold with a comparable reduction in colony output (Figures 6C and 6E). Colony output therefore does not scale with the number of reprogramming intermediates. Second, *TOX* knockdown also affected a program that defines reprogramming progression rather than proliferation: EMT and TGF-β signaling increased alongside the loss of proliferative programs (Figure 6G). Because MET is a required early step in fibroblast reprogramming and TGF-β signaling opposes it^25,26^, this shift is consistent with impaired acquisition or maintenance of the epithelial-like reprogramming program.

### *TOX* overexpression is not sufficient to enhance iPSC formation

Because *TOX* depletion impaired reprogramming, we next asked whether increasing *TOX* expression was sufficient to enhance it. We generated lentiviral Tet-On *TOX* HDFs and induced *TOX* during time windows corresponding to its endogenous transient expression (Figure 6H). Doxycycline treatment markedly increased *TOX* expression relative to Ctrl Dox (+), although cells transduced with the *TOX* overexpression vector showed a slight increase in *TOX* expression even without doxycycline (Figure 6I). *TOX* overexpression did not substantially improve iPSC colony formation under any induction schedule; colony output was numerically lower in several conditions, except under the no-doxycycline and day 10–14 doxycycline conditions (Figure 6J).

*TOX* induction produced few large changes at the individual-gene level beyond TOX itself (Figure S6G), but shifted the pathways affected by knockdown in the opposite direction: E2F target, G2M checkpoint, and MYC target programs increased, whereas EMT and TGF-β signaling decreased (Figure 6K). Thus, the overexpression construct was biologically active, yet it did not increase iPSC colony formation. Together with the loss-of-function results, these findings identify TOX as a context-dependent, permissive component of the OSKM network rather than a factor sufficient to drive iPSC formation. The shared loss-of-function phenotype, coupled with distinct gene-level responses, prompted us to examine the cell type-specific regulatory contexts in which TOX operates.

### Gene programs affected by *TOX* knockdown are predominantly cell type-specific

TOX was required in all three lineages, yet the genes it affected differed. We therefore determined what fraction of the TOX-dependent genes was shared between lineages and what fraction was lineage-specific. Pando nominates candidate target genes by linking a transcription-factor motif found in an accessible regulatory element to the expression of a nearby gene—a TF–CRE–target relationship.

Pando’s motif reference contains no TOX-specific motif; instead, its default motif-to-TF map^30^ assigns TOX three motifs of related HMG-box factors on the basis of protein-sequence similarity, which we used without modification (Figure S7A)^40,41^. These motifs are shared with other HMG-box factors, including the exogenous SOX2, so network inference alone cannot separate genes that depend on TOX from genes responding to another factor. We therefore required independent perturbational support: for each cell type we kept only the Pando-nominated genes that were also downregulated after *TOX* knockdown (Figure 7A), and we refer to these as KD-supported candidate genes.

**Figure 7.**
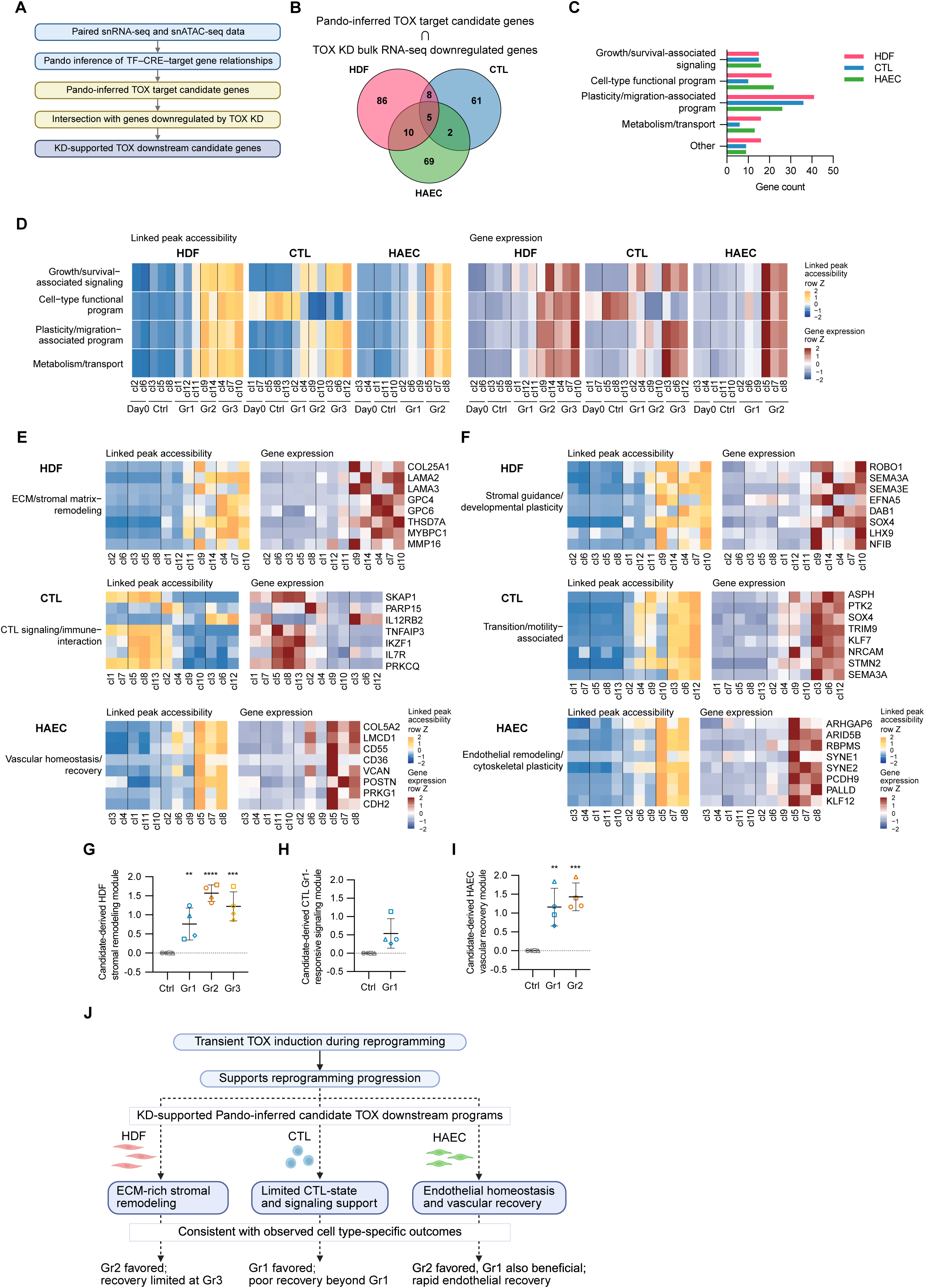
Shared TOX dependence is associated with predominantly lineage-specific candidate regulatory programs. (A) Schematic of the candidate-selection strategy. (B) Venn diagram showing overlap of KD-supported TOX downstream candidate genes across HDFs, CTLs, and HAECs. The filtered sets contained 109 HDF, 76 CTL, and 86 HAEC candidates, of which *COL25A1*, *HTR7*, *MAP7*, *PIR*, and *SYNE1* were shared across all three cell types. (C) Functional classification of KD-supported TOX downstream candidate genes in HDFs, CTLs, and HAECs. Pando-inferred TOX target candidates were intersected with genes that showed decreased expression after *TOX* knockdown in bulk RNA-seq. Categories include growth/survival-associated signaling, cell-type functional program, plasticity/migration-associated program, metabolism/transport, and other. (D) Category-level heatmaps of linked peak accessibility and gene expression across representative clusters; the “Other” category in panel C was not included. Values are row-scaled Z-scores calculated from category-level average linked peak accessibility or gene expression. The heatmaps show when each KD-supported TOX candidate program is most prominent along the reprogramming trajectory in each cell type. (E) Representative cell-type functional programs for each cell type. HDFs show an ECM/stromal matrix-remodeling program, CTLs show a CTL signaling/immune-interaction program, and HAECs show a vascular homeostasis/recovery program. For each module, linked peak accessibility and gene expression are shown as row-scaled Z-scores. (F) Representative plasticity/migration-associated programs for each cell type. HDFs show stromal guidance/developmental plasticity, CTLs show transition/motility-associated programs, and HAECs show endothelial remodeling/cytoskeletal plasticity. For each module, linked peak accessibility and gene expression are shown as row-scaled Z-scores. (G) Candidate-derived HDF stromal remodeling module score (11 genes) in HDFs at day 25, shown as paired condition-minus-Ctrl values. Statistical comparisons and displayed significance values were calculated from the corresponding original (uncentered) donor-level scores. The central horizontal line indicates the mean of four independent donors, and error bars indicate SD; open symbols indicate HDF donors circle 54F, square 48F, triangle 56M, and diamond 48M. Repeated-measures one-way ANOVA followed by Dunnett’s multiple comparisons test versus Ctrl was used. \*\**p* < 0.01, \*\*\**p* < 0.001, \*\*\*\**p* < 0.0001; unmarked comparisons are not significant. (H) Candidate-derived CTL Gr1-responsive signaling module score (6 genes) in CTLs at day 19, shown as paired condition-minus-Ctrl values. The statistical comparison was performed on the corresponding original (uncentered) donor-level scores using a two-tailed paired t test. The central horizontal line indicates the mean of four independent donors, and error bars indicate SD; open symbols indicate CTL donors circle 58M, square 55M, triangle 56F, and diamond 54F. No significant difference was detected. (I) Candidate-derived HAEC vascular recovery module score (10 genes) in HAECs at day 21, shown as paired condition-minus-Ctrl values. Statistical comparisons and displayed significance values were calculated from the corresponding original (uncentered) donor-level scores. The central horizontal line indicates the mean of four independent donors, and error bars indicate SD; open symbols indicate HAEC donors circle 63M, square 61M, triangle 51F, and diamond 50F. Repeated-measures one-way ANOVA followed by Dunnett’s multiple comparisons test versus Ctrl was used. \*\**p* < 0.01, \*\*\**p* < 0.001; unmarked comparisons are not significant. (J) Working model distinguishing experimentally supported relationships from inferred or phenotype-associated links. Transient *TOX* induction and loss-of-function data support a shared, permissive contribution to reprogramming progression across HDFs, CTLs, and HAECs. Integration of Pando inference with *TOX*-depletion transcriptomes nominates predominantly lineage-specific candidate programs involving stromal remodeling in HDFs, CTL-state signaling and cellular transition in CTLs, and endothelial homeostasis and vascular remodeling in HAECs. The state distributions and endpoint persistence of these programs parallel aspects of the distinct post-withdrawal phenotypes, with stronger endpoint support in HDFs and HAECs than in CTLs. Solid arrows denote experimentally supported relationships; dashed arrows denote inferred or phenotype-associated relationships. The model does not imply direct TOX binding or causal control of the recovery phenotypes.

Before KD-based filtering, Pando nominated 163, 185, and 219 candidate downstream genes of TOX in HDFs, CTLs, and HAECs, respectively, of which only 24 were shared across all three cell types (Figure S7B). Requiring KD support reduced the sets to 109, 76 and 86 candidates, and left only five genes— *COL25A1*, *HTR7*, *MAP7*, *PIR* and *SYNE1*—in common (Figure 7B). These five span matrix, signaling, cytoskeletal, nuclear-envelope and redox functions, and are the candidates for a lineage-independent component of TOX dependence. Because the phenotypes we set out to explain differed by lineage, we examined the biological programs represented within each lineage.

We next asked whether at least one of these candidate relationships held up at the locus level, and chose *COL25A1* because it passed KD-based filtering in all three cell types. Peak-to-gene analysis identified several candidate CREs linked to *COL25A1* across HDFs, CTLs, and HAECs (Figure S7C), and two representative elements were analyzed by state-resolved seq2PRINT sequence attribution and FIMO^42^ motif scanning (Figure S7D). In all three cell types both elements gained accessibility and attribution signal in more advanced reprogramming states, and both carried matches to the surrogate HMG-box motifs. *COL25A1* therefore shows the pattern expected of a gene regulated through *TOX*-linked elements during reprogramming: its elements open as cells progress, they carry HMG-box motif matches, and the gene falls when *TOX* is depleted. Motif matches cannot demonstrate binding, so *COL25A1* remains a candidate direct target rather than a validated one.

We next classified the broader sets of KD-supported candidates into four functional categories: growth/survival-associated signaling, cell type–functional programs, plasticity/migration-associated programs, and metabolism/transport (Figure 7C). HDF candidates were most frequently assigned to cell type–functional and plasticity/migration-associated programs. CTLs had the fewest candidates and showed a relative predominance of plasticity/migration-associated programs. HAECs showed a distribution broadly similar to that of HDFs, but with greater representation of cell type–functional programs.

If these candidate programs contribute to what cells retain after OSKM withdrawal, their activity should track the states that differed in recovery. We therefore mapped them across the reprogramming states of each cell type (Figure 7D). In HDFs and HAECs, all four categories rose modestly in Gr1 and then sharply in the deeper states, in both accessibility and expression. CTLs behaved differently: the cell type–functional program was highest in Ctrl and Gr1, whereas the other three categories peaked in Gr3. The same functional categories therefore peaked at different depths in each lineage—in HDFs and HAECs at the depths that retained a reduced DNA methylation age, and in CTLs at the depth from which recovery failed, except the cell type–functional program, which was highest in the states that recovered.

To ask whether the cell type–functional candidates matched the phenotypes we had measured, we examined their constituent genes (Figure 7E). In HDFs they formed a collagen-, laminin-and proteoglycan-rich ECM/stromal module^43–46^ that opened and rose in the deeper states, matching the stromal bias seen at the endpoint. In CTLs they formed a signaling and immune-interaction module (*SKAP1*, *IL12RB2*, *IKZF1*, *IL7R*, *PRKCQ*, *TNFAIP3*, *PARP15*)^47,48^ that was expressed mainly in day 0, Ctrl and Gr1 clusters and largely lost in Gr2/Gr3. In HAECs they comprised matrix-interaction, adhesion and vascular-signaling genes^49–51^, matching established programs of endothelial homeostasis and repair. In each lineage, then, the candidates named the very functions that did or did not come back. We examined the largest category, the plasticity/migration-associated candidates, in the same way (Figure 7F). These were semaphorin, Slit/Robo, ephrin and developmental regulators in HDFs^52–54^, adhesion, cytoskeletal and motility genes in CTLs^55,56^, and cytoskeletal, nuclear-architecture and adhesion genes in HAECs^49,57^; in every case they rose toward the deeper states, consistent with the remodeling that accompanies reprogramming progression.

The two remaining categories followed lineage-appropriate patterns: stromal growth-factor and niche signaling in HDFs^43,44^, adaptive receptor and kinase signaling in CTLs, vascular growth and survival signaling in HAECs^49–51^, and lipid, solute-transport and redox programs in all three, but aligned less directly with the measured phenotypes (Figures S7E and S7F).

Finally, we asked whether the phenotype-aligned subsets of these candidates were still elevated at the recovery endpoint. We built endpoint modules from the programs in Figures 7D–7F, keeping the genes detected in the matching endpoint bulk RNA-seq. In HDFs, an 11-gene stromal-remodeling module was significantly elevated relative to Ctrl in all three states at day 25 and highest in Gr2 (Figure 7G), matching the persistent stromal remodeling we measured. In HAECs, a 10-gene vascular-recovery module was significantly elevated in Gr1 and Gr2 at day 21 and highest in Gr2 (Figure 7I), matching the identity recovery and improved tube formation. In CTLs, the corresponding six-gene module rose only modestly and was not significantly elevated at day 19 (Figure 7H).

Together, these analyses support a model combining shared and lineage-specific features of human somatic cell reprogramming (Figure 7J). TOX contributes permissively to progression in all three cell types, yet the candidate programs associated with that dependence are predominantly lineage- and state-specific, and in HDFs and HAECs they remain elevated after factor withdrawal alongside the phenotypes they resemble. These associations do not establish causality, but they place a shared dependency inside three different regulatory contexts, consistent with a single input producing three different post-withdrawal outcomes.

## Discussion

The central challenge in translating partial reprogramming is not simply to reverse molecular marks of aging, but to define how far to reprogram the cells so that they durably recover their source-lineage identity and function after OSKM withdrawal. Our findings suggest that the answer to this question depends on lineage or cell type, as human fibroblasts, cytotoxic T cells, and aortic endothelial cells displayed different abilities to recover from partial reprogramming in our in vitro assays. Importantly, we did not find a strict relationship between depth of reprogramming and functional recovery that would apply across cell types. This finding suggests that for therapeutic applications, partial reprogramming protocols should not be defined by exposure duration alone and must be paired with post-withdrawal assessment of recovery. All three lineages depended on the HMG-box factor TOX to progress efficiently through reprogramming. However, candidate *TOX*-linked downstream programs were predominantly lineage-specific and paralleled aspects of the lineage-specific recovery phenotypes. Overall, our findings demonstrate that the tradeoff between cell rejuvenation and recovery of somatic function differs between cell types, but reveal shared and cell-type-specific molecular entry points through which we may eventually optimize this tradeoff.

Our study adds to and complements previous investigations of partial reprogramming and post-withdrawal recovery. Previous studies established that transient reprogramming can modify age-associated features while preserving somatic identity^11,13^, and interrupted reprogramming allowed human fibroblasts^14^ and multiple murine cell types^16^ to reacquire their identity after factor withdrawal. Cross-lineage analyses of full reprogramming further identified shared late-stage features^18^ and cell-of-origin-dependent routes to pluripotency^19^. These studies, however, did not systematically compare cells at different intermediate states of reprogramming and across multiple primary human lineages for their ability to recover both source-lineage identity and specialized function after OSKM withdrawal. In vivo studies in mice have established that cyclic or transient OSK(M) expression can improve tissue and organismal phenotypes^11,12^, including endothelium-directed OSK delivery that attenuates endothelial-to-mesenchymal transition and lowers blood pressure in hypertensive mice^17^. However, such readouts are integrated at the level of the tissue or animal and cannot resolve how far an individual lineage has progressed or what it retains from its reprogramming once the factors are withdrawn. The two approaches are therefore complementary: organism-level studies establish whether a therapeutic benefit accrues, whereas the state-resolved human measurements reported here define the depth at which that benefit can be obtained without losing the source lineage.

Our findings challenge three assumptions implicit in how partial reprogramming is currently designed and interpreted. The first is that a given OSKM exposure leaves different cell types in a comparable state, and therefore allows comparable recovery. Under identical exposure conditions, however, the three lineages reached different states at the same time point (Figure 1D), so fixed-duration comparisons conflate time under OSKM with the state actually reached. By prospectively isolating marker-defined states ranked by progression within each lineage, we could separate intermediate-state biology from reprogramming kinetics. Even states matched by ordinal rank were not equivalent: HDF Gr2 retained a reduced DNA methylation age but recovered proliferation only partially, whereas HAEC Gr2 rapidly regained endothelial identity together with markedly improved vascular function.

A second expectation is that lineages which reprogram efficiently to iPSCs will also recover well from partial reprogramming. CTLs formed iPSCs roughly 150-fold more efficiently than HAECs, yet only the shallowest CTL state recovered, and it did not retain its DNA methylation age reduction. HAECs, the least iPSC-competent lineage, showed the most favorable combination of retained methylation-age reduction, identity recovery, and improved specialized function under the conditions tested here. Competence for full pluripotency therefore does not predict the depth from which a lineage can recover, or the function it displays afterwards.

The third expectation is that partial reprogramming resets molecular age and then lets cells return to a normal state. In HDFs, the states that retained the methylation-age reduction also retained ECM and contractile stromal programs and incomplete proliferative recovery, so the post-withdrawal outcome was a mixed phenotype rather than uniform improvement. In CTLs, the methylation-age reduction was lost altogether during recovery. A persistent clock shift can therefore accompany an incompletely normalized lineage state, and a fully recovered lineage identity can come with no retained clock benefit at all.

The fibroblast outcome deserves a closer look, because fibroblasts dominate the human partial-reprogramming literature. Recent work showed that OSKM exposure can suppress aging-associated mesenchymal drift and activated-myofibroblast signatures in human fibroblasts^58^. Our findings add the recovery-phase perspective: although EMT-and TGF-β-associated activity declined during active reprogramming, ECM and contractile programs re-emerged after withdrawal, consistent with established matrix-remodeling and myofibroblast-like states^43–46^. A reduced DNA methylation age can therefore coexist with an incompletely normalized lineage state.

The constraints behind the CTL and HAEC outcomes are already known ones: CD28 extinction accompanies late T-cell differentiation^59^, whereas endothelial function depends on angiogenic and mechanotransduction programs^49–51,57^. Fibroblasts, T cells and endothelial cells therefore differ not only in how deep they can be taken, but in which functions must return before recovery can be called complete.

From a practical standpoint, the three lineages point in different directions. For HDFs, no state combined a retained methylation-age reduction with full functional recovery, so dermal fibroblasts appear the least favorable of the three for partial-reprogramming-based rejuvenation. This is worth noting, because most human work to date has used this cell type. For CTLs, Gr2 and Gr3 failed to sustain expansion after OSKM suppression whereas Gr1 outgrew matched controls, which suggests that continued stimulation after withdrawal could enrich for recovering cells without prospective sorting. HAECs were the only lineage in which all three of the benefits we measured were retained together, and are the most promising of the three. Because these three lineages span mesenchymal, hematopoietic, and endothelial origins and none behaved like another, per-lineage determination is likely to be a general requirement rather than a property of these particular cells.

These results have direct implications for developing partial-reprogramming strategies. Optimization should not ask only how long OSKM should be expressed, but which state represents an acceptable stopping point for the intended cell type and application: a uniform schedule could take a sensitive lineage past its recoverable range while leaving a useful state unexploited in another. Stopping criteria should therefore be state-based, defined with lineage-appropriate live markers, and paired with post-withdrawal confirmation of factor clearance, stable identity, proliferative competence and specialized function. The deepest attainable state should not be presumed to be the best one. Likewise, factor withdrawal alone cannot establish safety, because clearance of exogenous expression does not by itself demonstrate genomic stability, absence of aberrant states, or lack of tumorigenic potential^10,60^. Our data do not establish clinical safety or superiority over iPSC-based approaches; rather, they define the measurements needed to evaluate whether a partial-reprogramming intervention has achieved a durable and functional somatic endpoint.

Mechanistic studies of full iPSC induction have identified regulators of discrete reprogramming transitions, including *Nr0b1* and *Etv5* in prospectively isolated early intermediates^61^ and *TOX4*, a distinct paralog of *TOX*, during early reprogramming^32^. Context-specific mediators of beneficial partial-reprogramming effects have also been identified, including *TET1/2* in retinal ganglion cells^12^ and *Top2a* in hepatocytes^62^. However, these studies left unanswered the question of whether different lineages rely on similar or different gene programs to progress through reprogramming. A simple expectation is that since OSKM can reprogram a variety of lineages, reprogramming must depend on at least some shared pathways between lineages.

Indeed we identified TOX as a shared reprogramming dependency. TOX emerged from the compact Pando-inferred regulatory module shared among all three lineages, was transiently induced in advanced intermediate states, and its transient expression was accompanied by the opening of *TOX*-linked candidate cis-regulatory elements. *TOX* depletion reduced the establishment or maintenance of reprogrammed populations and sharply impaired iPSC derivation in all three cell types without reducing early SeV-OSKM transcript abundance. Reprogramming requires cell division^38,39^, and *TOX* depletion attenuated proliferative programs, so part of this phenotype may reflect reduced proliferation of the intermediates themselves. *TOX* knockdown in the absence of reprogramming factors largely preserved cell yield, however, indicating that the deficit is confined mainly to cells undergoing OSKM-driven progression rather than a general requirement for TOX in proliferation. Conversely, increasing *TOX* expression in HDFs altered progression-associated pathways but did not enhance iPSC formation. These asymmetric loss-and gain-of-function phenotypes support a role for TOX as a permissive progression dependency rather than an independently sufficient reprogramming driver.

Shared TOX dependence did not produce a shared downstream program. Integrating Pando-inferred relationships with cell type-matched *TOX*-knockdown transcriptomes yielded predominantly lineage-specific candidate genes and pathways. These programs fell into established lineage-relevant domains: ECM/stromal and developmental-guidance programs in HDFs^43–46,52–54^, immune-signaling and adhesion/motility programs in CTLs^47,48,55,56^, and vascular-homeostasis and cytoskeletal/mechanotransduction programs in HAECs^49–51,57^. Moreover, expression of the HDF and HAEC candidate modules was still elevated after OSKM withdrawal, paralleling stromal remodeling and vascular recovery, respectively. The corresponding CTL endpoint module was not significantly elevated, limiting inference in that lineage. These relationships remain associative rather than causal. Further experiments will be needed to determine which downstream candidates are direct targets of TOX, and which ones determine the post-withdrawal phenotypes.

Two features of TOX biology help explain why a single shared dependency can act through largely different programs in each lineage. First, TOX is an architectural HMG-box protein predicted to recognize DNA structure with relatively low sequence specificity^31^, so it would be expected to act on whatever regulatory landscape it encounters; consistent with this, TOX perturbation can reshape transcriptional and chromatin-accessibility programs in CD8+ T cells^63^. Second, because TOX is best known as a driver of CD8+ T-cell exhaustion^63–66^, it is worth saying explicitly that TOX induction alone should not be interpreted as evidence of exhaustion. *TOX* expression in humans is not restricted to exhausted cells and is also found in polyfunctional effector-memory populations^67^. Canonical exhaustion-associated genes were not prominent among the CTL candidates, and in the recoverable Gr1 population the functional data point the other way: CD28 positivity and effector output both increased. More broadly, our findings argue that partial reprogramming is a family of lineage-conditioned transitions rather than a single reversible intermediate shared across cell types. Cells of different origin start from different epigenetic landscapes^68,69^, take different routes to pluripotency^19^, diversify early in reprogramming^33–35^, and converge only late^18^. The same permissive factor therefore acts on a different accessible landscape in each lineage, and leaves a different residue behind once OSKM is withdrawn.

TOX dependence therefore identifies regulatory convergence without erasing lineage divergence. TOX and its candidate downstream programs suggest three directions for optimizing partial reprogramming. First, induction of TOX and opening of shared candidate regulatory elements at the TOX locus point to a reporter strategy that could be tested across lineages, including cell types lacking established surface-marker panels, because TOX was induced in all three lineages whereas the marker gates used here had to be defined separately for each. Such reporters would require lineage-specific calibration against post-withdrawal recovery: high TOX in HAEC Gr2 coexisted with robust identity and functional recovery. Second, timed or titrated TOX suppression could test whether progression can be restricted to shallower states while retaining recoverable cells and their functional benefits. In CTLs, the key test would be whether these interventions preserve the recoverable Gr1 population, which was also reduced by knockdown. Third, the TOX-linked candidate modules give lineage-specific readouts for the recovery phase. In HDFs, the stromal module remained elevated above matched controls at the recovery endpoint, marking the residual matrix bias to be avoided; in HAECs, the vascular module tracked the functional recovery to be achieved. Panels of this kind could be measured alongside functional assays when protocols are compared. The same modules also define an experiment: in HDFs, perturbing selected genes during recovery would test whether persistent stromal activation can be reduced, and proliferative recovery improved, while the methylation-age reduction is retained. That would act on one unwanted component of the recovery response, rather than on TOX itself or on the matrix functions fibroblasts need.

### Limitations of the study

Our experiments were performed in vitro using primary cells from four middle-aged donors per cell type and therefore do not capture the full diversity of age, tissue, and multicellular contexts encountered in vivo. DNA methylation age estimates are clock-, cell type-, and culture-time-dependent and should be interpreted together with transcriptomic and functional readouts. KD support was defined directionally, by reduced expression after *TOX* depletion rather than by a statistical threshold, so individual candidates require independent confirmation. Finally, integration of computational inference with acute TOX perturbation supports TOX-associated regulatory relationships but does not establish direct TOX binding to the nominated CREs or causal control of the associated rejuvenation and functional phenotypes. Genome-wide TOX occupancy data remain limited and their interpretation is debated, consistent with the classification of TOX within the sequence-independent HMG-box subfamily^31,70^, so the cis-regulatory relationships reported here should be treated as candidate rather than validated targets.

## METHODS

### RESOURCE AVAILABILITY

#### Lead contact

Requests for further information and resources should be directed to and will be fulfilled by the lead contact, Akira Kunitomi.

#### Materials Availability

Requests for the customized SeV vectors should be directed to the lead contact, who can facilitate contact with the provider. Other unique reagents generated in this study are available from the lead contact under a Materials Transfer Agreement.

#### Data and Code Availability

The single-nucleus multiome, bulk RNA sequencing, and DNA methylation array datasets generated in this study have been deposited in the NCBI Gene Expression Omnibus and will be made publicly available upon publication of the peer-reviewed version of this study.

All custom code used for partial reprogramming and gene regulatory network analyses will be made publicly available through Zenodo and GitHub upon publication of the peer-reviewed article.

An interactive web-based browser built with the UCSC Cell Browser^71^ has been developed for exploring the single-nucleus multiome data, RNA velocity, chromatin-accessibility profiles and scPrinter/seq2PRINT footprints at the *TOX* and *COL25A1* loci, and the Pando-inferred gene regulatory networks. The browser will be made publicly available upon publication of the peer-reviewed article.

The trained scPrinter/seq2PRINT models for HDFs, CTLs, and HAECs, comprising the bulk sequence models and the corresponding stage-specific LoRA fine-tuned models, will be made publicly available through Hugging Face upon publication of the peer-reviewed article.

### EXPERIMENTAL MODEL AND STUDY PARTICIPANT DETAILS

Primary HDFs (NHDF-Ad-Der Fibroblasts FGM-2, Lonza, Cat# CC-2511), primary HAECs (HAEC-Aortic Endothelial EGM-2, Lonza, Cat# CC-2535), and CD8+ CTLs isolated from commercially obtained human PBMCs (Cellular Technology Ltd., identifier CTL-UP1) were used. Four independent donors were analyzed for each cell type; donor age and sex are indicated in the corresponding figure legends. Before SeV infection, HDFs were maintained at 37°C under 5% CO_2_ and 2% O_2_, whereas HAECs and CTLs were maintained at 37°C under 5% CO_2_ and 21% O_2_. Detailed culture, SeV transduction, temperature-shift, and cell-sorting procedures are described below. All work involving recombinant DNA and recombinant Sendai virus vectors was conducted under protocols approved by the Institutional Biosafety Committee of Gladstone Institutes.

### METHOD DETAILS

#### Human primary cell culture

Normal human dermal fibroblasts (HDFs; NHDF-Ad-Der; Lonza) were cultured in high-glucose DMEM containing GlutaMAX Supplement (Gibco), supplemented with 10% heat-inactivated fetal bovine serum (Sigma-Aldrich), 1× MEM non-essential amino acids (Gibco), and 50 µM 2-mercaptoethanol (Gibco).

Cells were maintained on gelatin-coated culture dishes (STEMCELL Technologies). Medium was replaced every other day, and cells were passaged using 0.05% Trypsin-EDTA (Gibco) every 5–7 days, depending on confluence.

Human CD8+ cytotoxic T lymphocytes (CTLs) were isolated from peripheral blood mononuclear cells (Cellular Technology Ltd.) using the EasySep Human CD8+ T Cell Isolation Kit and EasySep Buffer (STEMCELL Technologies). CTLs were cultured in X-VIVO 15 medium (Lonza) supplemented with 5% human serum (Sigma-Aldrich), 10 mM N-acetyl-L-cysteine (Sigma-Aldrich), 5 ng/mL recombinant human IL-7 (PeproTech), and 5 ng/mL recombinant human IL-15 (PeproTech). For routine maintenance, cells were collected every 7 days, the medium was replaced, and the cells were stimulated with Dynabeads Human T-Activator CD3/CD28 (Gibco) at 25 µL per 1 × 10^6^ cells. An equal volume of fresh medium was added 2 days after stimulation. Dynabeads were removed and the medium was replaced on day 4, followed by the addition of an equal volume of fresh medium on day 6.

Human aortic endothelial cells (HAECs; HAEC-Aortic Endothelial; Lonza) were cultured in EBM Endothelial Cell Growth Basal Medium (Lonza) supplemented with the EGM-2 Endothelial SingleQuots Kit (Lonza). Medium was replaced every other day, and cells were passaged using 0.05% Trypsin-EDTA (Gibco) every 4–6 days, depending on confluence.

#### Reprogramming of human somatic cells using temperature-sensitive Sendai virus vectors

HDFs and HAECs were seeded at 2 × 10^4^ cells/cm² 1 day before SeV transduction (day −1). CTLs were stimulated on day −4 with Dynabeads Human T-Activator CD3/CD28 (Gibco) at 75 µL per 1 × 10^6^ cells. The day of SeV transduction was defined as day 0.

On day 0, CTLs were transferred to V-bottom 96-well plates at 5 × 10^5^ cells in 100 µL per well. HDFs, CTLs, and HAECs were transduced with the following three temperature-sensitive SeV vectors added simultaneously to the culture medium:

SeV(PM)KOS-DGFP/TS12ΔF (ID Pharma), multiplicity of infection (MOI) 5
SeV(HNL)C-MYC-OFP/TS15ΔF (ID Pharma), MOI 3
SeV18+KLF4/TS12ΔF (ID Pharma), MOI 5

Control cells were transduced with the corresponding temperature-sensitive fluorescent control-vector set lacking the reprogramming-factor cassettes: SeV18+EGFP/TS15ΔF, SeV(HNL)OFP/TS15ΔF, and SeV/TS15ΔF (all from ID Pharma), at the corresponding MOIs of 5, 3, and 5, respectively. Aliquots of uninfected cells collected before transduction were used as day 0 samples.

To improve transduction efficiency, CTLs were subjected to spinoculation at 1,000 × g and 22°C for 30 min immediately after addition of the SeV vectors. To maintain expression from the temperature-sensitive SeV vectors, cells were cultured at 35°C in a humidified CO_2_ incubator. HDFs were maintained at 2% O_2_. The medium was replaced with the corresponding cell type-specific medium on the following day and every other day thereafter.

One day after SeV transduction, CTLs were transferred to 12-well plates coated with iMatrix-511 silk (Matrixome) at up to 1 × 10^6^ cells per well. The medium was changed to MSC NutriStem XF Basal Medium (Sartorius), and the cells were maintained under adherent culture conditions. HDFs and HAECs were harvested on day 7, and CTLs were harvested on day 5, for flow cytometric analysis and cell sorting.

#### Flow cytometric analysis and cell sorting

HDFs and HAECs were dissociated into single-cell suspensions on day 7 using 0.05% Trypsin-EDTA (Gibco). CTLs were dissociated on day 5 using TrypLE Select (Gibco). Cells were resuspended in FACS buffer consisting of PBS, 2% fetal bovine serum (Sigma-Aldrich), and 2 mM EDTA (IBI Scientific), and stained for 15 min on ice in the dark with fluorophore-conjugated antibodies diluted 1:20. Zombie Violet Fixable Viability Kit or Zombie R718 Fixable Viability Kit (BioLegend) was used to exclude dead cells.

After staining, cells were washed with FACS buffer and resuspended in the appropriate phenol red-free medium supplemented with Antibiotic-Antimycotic (Gibco) and 2 mM EDTA: DMEM, high glucose, no glutamine, no phenol red (Gibco) for HDFs; TheraPEAK X-VIVO 15 (Lonza) for CTLs; and EBM Endothelial Cell Growth Basal Medium, Phenol Red Free (Lonza) for HAECs. Cells were analyzed using a Cytek Aurora (Cytek Biosciences) and sorted using a BD FACSAria Fusion (BD Biosciences). After exclusion of doublets and dead cells, cells positive for both GFP and OFP were operationally defined as the SeV-OSKM-enriched population. GFP+OFP+ cells were further classified into Gr1, Gr2, and Gr3 according to the expression of lineage-associated and early reprogramming markers: CD13 and SSEA5 for HDFs, CD8α and SSEA4 for CTLs, and CD31 and SSEA4 for HAECs. Flow cytometry data were analyzed using FlowJo v10.10.0 (FlowJo, LLC).

Immediately after sorting, aliquots of each population were collected for bulk RNA-seq and DNA methylation array analyses. The remaining cells were used for temperature-shift-mediated termination of SeV-OSKM expression and post-sort recovery assays.

#### Temperature-shift-mediated termination of SeV-OSKM expression and post-sort recovery culture

Immediately after sorting, the culture temperature was shifted from 35°C to 38°C to suppress replication of the temperature-sensitive SeV genomes and terminate SeV-derived reprogramming-factor expression. HDFs were maintained on iMatrix-511 silk-coated culture dishes until day 14 and were transferred back to gelatin-coated dishes at the passage on day 14. HAECs were maintained on gelatin-coated dishes throughout post-sort culture. HDFs and HAECs were passaged approximately once per week and maintained until the indicated endpoint analyses, primarily day 25 for HDFs and day 21 for HAECs.

After sorting, CTLs were returned to suspension culture in the T-cell medium described above and maintained at 38°C. CTLs were restimulated weekly using Dynabeads Human T-Activator CD3/CD28 (Gibco) and maintained until day 19 or the indicated later analysis time points.

#### iPSC colony formation assay

iPSC colonies were counted on day 14 after SeV infection. Cells were fixed and stained using the StemAb Alkaline Phosphatase Staining Kit II (ReproCell) according to the manufacturer’s protocol, and AP-positive colonies were counted using an Olympus CKX41 microscope. Reprogramming efficiency was calculated as the number of AP-positive iPSC-like colonies divided by the number of cells seeded at the final plating step, multiplied by 100.

#### Bulk RNA sequencing and data processing

Total RNA and DNA were extracted from cell lysates using the AllPrep DNA/RNA Mini Kit or AllPrep DNA/RNA Micro Kit (QIAGEN), and RNA samples were treated with RNase-Free DNase Set (QIAGEN) to remove genomic DNA. Extracted RNA was submitted to Novogene or Plasmidsaurus for library preparation and bulk RNA sequencing. Quality control, read processing, alignment to the human reference genome, and gene-level quantification were performed by the respective service providers using their standard pipelines. Novogene-generated gene-level raw count matrices were used for the HDF, CTL, and HAEC reprogramming/follow-up datasets, whereas Plasmidsaurus-generated differential expression tables were used for the *TOX* knockdown datasets unless otherwise indicated.

For Novogene raw-count datasets, lowly expressed genes were filtered by retaining genes with counts ≥10 in at least four samples, except for CTL datasets with fewer samples per condition, for which genes with counts ≥10 in at least two samples were retained. Differential expression analysis and variance-stabilizing transformation were performed in R using DESeq2^72^ with a donor-paired design (∼ donor + group). Contrasts were specified within each cell type and time point to compare reprogramming conditions with the matched Ctrl group. *p* values were adjusted using the Benjamini-Hochberg method. For Plasmidsaurus *TOX* knockdown datasets, vendor-generated differential expression tables comparing *TOX* siRNA/shRNA-treated samples with matched negative or scramble controls were used as provided. For Plasmidsaurus-derived datasets analyzed from count or CPM tables, differential expression was performed using limma/voom or limma with donor included in the model.

Preranked gene set enrichment analysis was performed using MSigDB^73,74^ gene sets with fgsea^75^ in R. DESeq2-based analyses used the Wald statistic as the ranking metric, limma-based analyses used the moderated t statistic, and vendor-generated Plasmidsaurus DGE tables were ranked by log2 fold change. When multiple rows mapped to the same gene symbol, the row with the largest absolute ranking statistic or log2 fold change was retained. For volcano-plot visualization, genes with FDR < 0.05 and an absolute log2 fold change > 1 were highlighted as differentially expressed.

For custom transcriptomic module-score analyses, predefined gene sets listed in Table S1 were intersected with genes detected in each expression matrix after filtering and gene-symbol mapping. Module scores were calculated as the arithmetic mean of transformed expression values for the detected genes in each gene set. For count-matrix datasets, transformed expression values were DESeq2 variance-stabilized values or gene-wise Z-scores calculated from log2(size-factor-normalized count + 1), depending on the analysis. TOX-associated endpoint modules (Figures 7G–7I) were restricted to KD-supported Pando-inferred candidates and scored using gene-wise Z-scores. Plotted module scores for paired datasets were centered to the matched donor Ctrl value by subtracting the mean Ctrl score within each donor.

#### Genome-wide DNA methylation profiling and DNA methylation age estimation

Bisulfite conversion of 500 ng genomic DNA was performed using the EZ-96 DNA Methylation-Lightning MagPrep kit (Zymo Research), and genome-wide DNA methylation was profiled using the Infinium MethylationEPIC BeadChip Kit (Illumina) according to the manufacturer’s protocols. DNA methylation β values were exported from GenomeStudio V2011.1 as FinalReport files. Downstream DNA methylation age analyses were performed directly from the exported AVG_Beta values; no additional background correction, probe-type normalization, or minfi-based preprocessing was applied after GenomeStudio export. Sample-level β-value distributions and clock-CpG coverage were inspected before age estimation.

CpG identifiers were matched to each clock model by removing suffixes from GenomeStudio TargetIDs to obtain cg identifiers. Replicate probe entries mapping to the same CpG were collapsed by averaging β values. DNA methylation age was estimated using the Horvath multi-tissue clock^4^ and the BLUP clock^27^ implemented in the methylclock^76^ R package, with coefficient data provided by methylclockData and with normalization disabled.

IntrinClock^28^ age was calculated using the model and code obtained from the authors’ Zenodo repository. For IntrinClock, untransformed GenomeStudio AVG_Beta values were extracted for model CpGs; missing model CpGs were imputed as β = 0.5 following the downloaded IntrinClock example workflow.

#### Quantitative PCR

For qPCR, 500 ng of DNase-treated total RNA was reverse-transcribed using PrimeScript RT Master Mix (Takara Bio), which contains oligo(dT) primers and random hexamers. qPCR was performed on a StepOnePlus instrument (Applied Biosystems) or LightCycler 96 instrument (Roche) using TaqMan Fast Advanced Master Mix (Applied Biosystems) according to the manufacturers’ instructions. Target transcripts and *GAPDH* were quantified in duplex reactions using FAM-and VIC-labeled TaqMan assays, respectively. Relative expression was calculated using the 2^−ΔΔCt^ method, with *GAPDH* as the endogenous reference and the corresponding control condition as the calibrator.

#### Scratch wound closure

HDFs were seeded in 24-well plates at 8 × 10^4^ cells/well. On the following day (day 0), cells were incubated for 2 h in DMEM, high glucose, GlutaMAX Supplement (Gibco) containing 1% FBS (Sigma-Aldrich). After reaching confluence, a linear scratch was generated using a sterile P1000 pipette tip. Detached cells were removed by washing three times with DPBS (Gibco), followed by replacement with fresh medium. Phase-contrast images of the same fields were acquired immediately after scratching (0 h) and 6 h later using an Olympus microscope. Wound width was measured at identical positions, and scratch closure was calculated as: scratch closure (%) = (W0 − W6) / W0 × 100, where W0 and W6 denote wound width at 0 h and 6 h, respectively.

#### Multiplex analysis of CTL-secreted factors

For endpoint cytokine analysis, CTLs were restimulated with Dynabeads Human T-Activator CD3/CD28 on day 26 after SeV-OSKM transduction, and culture supernatants were collected four days later on day 30. Supernatants were clarified by centrifugation to remove cellular debris and stored frozen until analysis. Secreted factors were quantified using the LEGENDplex Human CD8/NK Panel (13-plex) with V-bottom Plate, version 2 (BioLegend), according to the manufacturer’s instructions. Samples were analyzed without dilution and measured in technical triplicate. Fluorescence data were acquired using an Attune NxT Flow Cytometer (Thermo Fisher Scientific), and analyte concentrations were calculated using the LEGENDplex Data Analysis Software (BioLegend).

#### Tube formation assay

A 48-well plate was coated with 50 µL per well of growth factor-reduced Matrigel (Corning) and incubated at 37°C for 1 h. HAECs were seeded onto Matrigel-coated wells at 2 × 10^4^ cells per well in 100 µL of medium. After 4 h, three non-overlapping phase-contrast fields per well were imaged using an Olympus CKX41 microscope. To estimate cell number, PrestoBlue Cell Viability Reagent (Thermo Fisher Scientific) was added according to the manufacturer’s instructions, and fluorescence was measured using a SpectraMax i3x plate reader (Molecular Devices) at 560-nm excitation and 590-nm emission. Cell numbers were estimated using a pre-established standard curve, and each tube-formation metric was normalized to the estimated cell number in the corresponding well. Tube formation was quantified using the Angiogenesis Analyzer plugin^77^ in Fiji v1.54p^78^ by measuring the numbers of junctions, segments, and master segments and the total master segment length. Three technical replicate wells were analyzed per condition, and the mean of three fields was calculated for each well.

#### Single-nucleus multiome library preparation, sequencing, and primary processing

Nuclei were isolated from HDFs and HAECs according to the 10x Genomics protocol CG000366, Rev D, and from CTLs according to protocol CG000365, Rev B. Paired single-nucleus gene expression and chromatin accessibility libraries were generated using the Chromium Next GEM Single Cell Multiome ATAC + Gene Expression platform (10x Genomics) according to the manufacturer’s instructions, with a targeted recovery of 5,000 nuclei per sample.

Libraries were sequenced by the UCSF Center for Advanced Technology using an Illumina NovaSeq X Plus. ATAC libraries were sequenced using a 10B flow cell with a 51 × 12 × 24 × 51-cycle configuration for Read 1, i7, i5, and Read 2, respectively. Gene expression libraries were sequenced using a 25B flow cell with a 28 × 10 × 10 × 90-cycle configuration.

Demultiplexed FASTQ files were processed using Cell Ranger ARC v2.0.2 (10x Genomics) with the cellranger-arc count pipeline and the refdata-cellranger-arc-GRCh38-2024-A reference package. The resulting filtered gene expression and chromatin accessibility matrices, peak calls, ATAC fragment files, and associated cell-level metadata were used for downstream analyses.

#### Single-nucleus multiome processing and quality control

Single-nucleus multiome datasets from HDFs, CTLs, and HAECs were processed separately using ArchR^79^ with the human hg38 reference genome. Gene annotations derived from EnsDb.Hsapiens.v86 were used for RNA count-matrix import and downstream gene-centric analyses. Per-sample ATAC fragment files were converted into ArchR Arrow files using createArrowFiles(), with generation of a 500-bp TileMatrix and a GeneScoreMatrix. Doublet scores were calculated using addDoubletScores(), and predicted doublets were removed using filterDoublets().

Arrow files were constructed from nuclei with ≥1,000 ATAC fragments and a TSS enrichment score ≥4. Nuclei were then retained if they had ≥6,000 ATAC fragments, a TSS enrichment score ≥5, >2,000 RNA UMIs, >1,000 detected genes, and a mitochondrial transcript fraction <25%. Nuclei without matched RNA and ATAC profiles were excluded. All downstream analyses used the resulting cell type– specific quality-controlled nuclei: 78,269 HDF, 71,382 CTL and 48,202 HAEC nuclei.

#### ATAC dimensionality reduction and batch correction

ATAC dimensionality reduction was performed on the TileMatrix using addIterativeLSI(). CTLs and HAECs were processed using iterations = 2, LSIMethod = 2, varFeatures = 15,000, outlierQuantiles = c(0.06, 0.982), and dimsToUse = 1:16. HDFs were processed using iterations = 1 and LSIMethod = 1, with the remaining addIterativeLSI() parameters left at their ArchR defaults. Sample-associated variation in the LSI representation was corrected using addHarmony()^80^ with groupBy = “Batch” in all three cell types. For CTLs, clustering and UMAP embedding used the uncorrected iterative LSI representation, whereas the Harmony-corrected representation was used for HDFs and HAECs and for peak-to-gene linkage in all three cell types.

Final UMAP embeddings were generated using addUMAP(). CTLs used dimensions 3–16 of the uncorrected iterative LSI representation with nNeighbors = 30, minDist = 0.20, and metric = “cosine.” HDFs used dimensions 2–16 of the Harmony-corrected representation with nNeighbors = 20, minDist = 0.08, and metric = “cosine.” HAECs used dimensions 3–16 of the Harmony-corrected representation with nNeighbors = 30, minDist = 0.15, and metric = “cosine.”

#### Leiden clustering

Leiden clustering was performed separately for CTL, HDF, and HAEC using Scanpy^81,82^ and 20-nearest-neighbor graphs. The graphs were constructed from LSI dimensions 3–16 for CTL, the exported Harmony representation for HDF, and Harmony dimensions 3–16 for HAEC. The Leiden resolution parameters were set to 1.0, 1.2, and 0.8 for CTL, HDF, and HAEC, respectively. Cluster assignments and UMAP coordinates were transferred to ArchR by exact cell-barcode matching for downstream analysis and visualization.

#### Peak calling

Peak calling was performed independently for CTL, HDF, and HAEC datasets. Pseudo-bulk coverages were generated for each final Leiden cluster using ArchR addGroupCoverages(), and reproducible peak sets were called with MACS2^83^ through addReproduciblePeakSet(). Peak calling was grouped by leiden_10, leiden_12, and leiden_08 for CTL, HDF, and HAEC, respectively. Cell-by-peak matrices and GC-and accessibility-matched background peaks were generated using addPeakMatrix() and addBgdPeaks(). Peak sets were retained independently for each dataset and were not merged across cell types.

#### Peak-to-gene linkage analysis

Peak-to-gene (P2G) links were inferred independently for HDFs, CTLs, and HAECs using the ArchR function addPeak2GeneLinks(). Peak accessibility from the PeakMatrix was correlated with gene expression from the GeneExpressionMatrix across cell aggregates constructed using the Harmony-corrected ATAC representation in all three cell types. Candidate peak–gene pairs within 550 kb were evaluated. Links were retrieved using getPeak2GeneLinks() with resolution = 1 and retained using correlation cutoffs of 0.35 for HDFs and CTLs and 0.50 for HAECs, together with FDRCutOff = 1 × 10−4, varCutOffATAC = 0.25, and varCutOffRNA = 0.25.

For long-range analysis at the *TOX* locus, addPeak2GeneLinks() was rerun in each cell type–specific ArchR project after peak calling, with a maximum distance of 1.5 Mb. Links assigned to *TOX* were retrieved using a correlation cutoff of 0.20 and the same FDR and variance-quantile filters described above. For promoter-centered visualization, loop objects were restricted to arcs intersecting a ±2-kb window centered on the *TOX* transcription start site, and pyGenomeTracks arcs were drawn from this anchor to the retained *TOX*-linked peaks. Peaks and arcs overlapping the neighboring *NSMAF* promoter (chr8:58,659,006–58,660,443; hg38) were excluded. Regions were classified as shared only when independently called peaks overlapped in all three datasets. P2G correlations were used as link scores, and FDR bins were used to scale arc widths.

For the *COL25A1* locus, P2G tables generated using the standard settings described above were filtered for links assigned to *COL25A1* within chr4:108,753,053–109,443,814 (hg38). Linked peaks were connected to the *COL25A1* transcription start site and visualized together with cell type–specific chromatin-accessibility tracks using pyGenomeTracks^84^.

#### Genomic locus visualization

Genomic locus plots were generated on the GRCh38 (hg38) assembly using pyGenomeTracks. Preprocessed chromatin-accessibility bigWig tracks were referenced through saved Integrative Genomics Viewer (IGV) sessions, which defined the initial track set and display settings, including track order, colors, heights, and highlighted intervals. Custom Python scripts parsed the IGV session XML and generated pyGenomeTracks configuration files. During conversion, track names were simplified, dataset order was standardized, and y-axis ranges were harmonized across comparable tracks. Peak-to-gene link tracks generated as described above were incorporated into the corresponding configurations. Gene annotations were supplied from local files. The *TOX* locus was annotated using a GTF file, whereas the *COL25A1* locus was annotated using the UCSC ncbiRefSeqSelect table converted to BED12 format. Plotting intervals were either specified directly in locus-specific scripts or derived from the target gene and neighboring promoters. For *COL25A1*, the displayed interval was extended from the gene body to the nearest upstream and downstream promoters, with an additional 10-kb margin at both boundaries.

Regulatory regions of interest were displayed as vertical highlights using BED coordinates. At the *TOX* locus, intervals were obtained from a curated table and converted to centered BED regions. At the *COL25A1* locus, coordinates of the highlighted candidate CREs were recovered from the corresponding IGV session. When region labels were required, a separate BED annotation track was added because the native pyGenomeTracks highlight overlay does not display region names.

#### Cluster-level differential gene-expression analysis

Differential gene expression was evaluated separately within CTL, HDF, and HAEC using all final quality-controlled cells. The ArchR GeneExpressionMatrix was transferred to RNA-only Seurat objects using Seurat^85^ and normalized using LogNormalize with a scale factor of 10,000. Cell identities were set to the final Leiden clusters (leiden_10 for CTL, leiden_12 for HDF, and leiden_08 for HAEC). Seurat FindAllMarkers() was used to compare each cluster with all remaining cells using a two-sided Wilcoxon rank-sum test with only.pos = FALSE, min.pct = 0.1, and logfc.threshold = 0.01. Bonferroni-adjusted *p* values reported by Seurat were used. For cluster-level pathway analysis, genes were ranked by average log2 fold change for each representative cluster, and preranked GSEA was performed using MSigDB Hallmark gene sets with fgsea.

#### Joint HDF, CTL, and HAEC RNA and ATAC embeddings

Gene-expression count matrices were obtained from Cell Ranger-formatted filtered_feature_bc_matrix.h5 files and analyzed using Scanpy and rapids-singlecell. Cells were initially retained if they had fewer than 10,000 detected genes and <20% mitochondrial counts, and final inclusion was restricted to curated cell IDs that had passed the preceding RNA and ATAC quality-control workflow in ArchR. Counts were normalized to 10,000 counts per cell and log-transformed, while raw counts were retained in the analysis object. The 5,000 most highly variable genes were used for dimensionality reduction, with *POU5F1*, *SOX2*, *KLF4*, and *MYC* manually included when present. Total counts and mitochondrial percentage were regressed out, followed by scaling and PCA.

Harmony correction was performed separately within HDF, CTL, and HAEC using donor identity as the batch variable. The corrected embeddings from all three cell types were then combined, and a shared nearest-neighbor graph was constructed from the first 30 Harmony-corrected principal components using 350 neighbors. UMAP was generated with min_dist = 0.5, spread = 1.2, spectral initialization, and random seed 0.

#### Joint chromatin-accessibility embedding

For the corresponding scATAC-seq analysis, Cell Ranger fragment files and peak-count matrices aligned to hg38 were analyzed using Signac after restriction to the same ArchR-curated cell IDs. Accessibility was quantified over 5,553 consensus regions derived from aggregate accessibility across Cell Ranger peak matrices. This small set of regions, shared across cell types, was used only to place HDF, CTL, and HAEC cells in a common embedding. Peak counts were normalized by term frequency-inverse document frequency and decomposed into 50 latent semantic indexing components. The first LSI component was omitted from downstream analysis.

Harmony correction was performed separately within HDF, CTL, and HAEC on LSI components 2–30 using donor identity as the batch variable. For the final joint representation, the first 10 centered within-population Harmony dimensions were added to population centroids derived from uncorrected LSI components 2–11. These centroids were displaced 1.5-fold from the global centroid to preserve separation among HDF, CTL, and HAEC, and the resulting dimensions were standardized. UMAP was generated from this representation using cosine distance, 1,000 neighbors, min_dist = 0.35, spread = 2.5, and random seed 0. This joint embedding was used only to display chromVAR motif deviations in Figure S7A and was not used for any quantitative inference.

#### UniTVelo RNA velocity and latent time

RNA velocity was inferred independently for HDF, CTL, and HAEC using UniTVelo^29^ and cell type-matched spliced and unspliced RNA matrices. Only cells shared between the annotated single-nucleus dataset and the corresponding loom files were retained. Cell annotations, Harmony-corrected RNA principal components, and global RNA UMAP coordinates were transferred to the cell type-specific AnnData objects. A nearest-neighbor graph was constructed in Scanpy using the first 30 Harmony-corrected principal components and 30 neighbors.

UniTVelo was run using the precomputed neighbor graph, with the UMAP embedding as the basis, unified-time fitting (FIT_OPTION = “1”), AGENES_R2 = 1, and a maximum of 2,000 iterations. Cells in d7g1 were specified as roots for HDF and HAEC, whereas d5g1 cells were specified as roots for CTL (stage labels denote the sampling day followed by c for Ctrl or g1–g3 for Gr1–Gr3, so that d7g1 denotes the day 7 Gr1 population). For CTL, the leftmost d5g1 cell along the global RNA UMAP1 axis was used as a deterministic root anchor, and NUM_REP=2 was used to stabilize trajectory directionality.

Following model fitting, scVelo was used to calculate the velocity graph and project velocity vectors onto the global RNA UMAP^86^. For the combined visualization, cell type-specific UMAP-projected velocity vectors were matched to cells in the full three-cell-type UMAP by cell ID. The vectors were smoothed over a regular two-dimensional grid, with extreme velocity magnitudes and low-density regions masked, and visualized as streamlines using Matplotlib.

#### chromVAR motif-deviation analysis for integrated ATAC UMAP visualization

For chromVAR^40^ analysis in Figure S7A, reproducible peak sets previously generated separately for HDF, CTL, and HAEC in ArchR were exported, combined, and merged with BEDTools^87^ (merge -d 2000), yielding a unified set of 244,828 regions. Accessibility within these regions was quantified separately for each sample using SnapATAC2^88^. Motif occurrences were annotated using the Pando-provided motif matrices (see Pando GRN inference), and bias-corrected chromVAR deviation Z-scores were calculated using the GPU-enabled Python implementation in scPrinter with 50 background peaks matched for GC content and aggregate accessibility. The analysis focused on the three HMG-box motif models that Pando assigns to TOX on the basis of protein-sequence similarity: MA0077.1 (SOX9-like), MA0442.2 (SOX10-like), and MA0523.1 (TCF7L2-like). Per-cell deviation scores were combined across samples and projected onto the fixed all-cell ATAC UMAP. chromVAR scores were used only for coloring the embedding and were not used to construct the UMAP.

#### Pando GRN inference

Gene regulatory networks were inferred independently for the CTL, HDF, and HAEC single-nucleus multiome datasets using Pando v1.1.1^30^. The analyses used the quality-controlled nuclei defined above. For each dataset, RNA and chromatin-accessibility counts were extracted from the corresponding RNA-integrated ArchR project using the GeneExpressionMatrix and PeakMatrix and converted into matched Seurat/Signac objects. Peaks detected in at least one cell were retained in a Signac ChromatinAssay, and gene annotations were obtained from EnsDb.Hsapiens.v86 with UCSC-style hg38 coordinates.

Pando objects were initialized from the RNA and peak assays using the phastConsElements20Mammals.UCSC.hg38 conserved regions supplied with Pando. Candidate regulatory regions were defined by intersecting conserved elements with accessible peaks and excluding exonic sequence. Motif scanning was performed using the extended motif resource distributed with Pando v1.1.1. This resource contains 1,590 position-frequency matrices, including 1,024 derived from JASPAR 2020^41^ and 566 from CIS-BP^89^, together with sequence-similarity-based motif-to-TF assignments. The Pando-provided motifs and motif2tf resources were supplied to find_motifs() to scan candidate regulatory regions against the hg38 genome (BSgenome.Hsapiens.UCSC.hg38). RNA counts were log-normalized in Seurat with a scale factor of 10,000, and 2,000 variable genes were selected independently for each dataset using FindVariableFeatures(selection.method = “vst”); no genes were force-added. Pando infer_grn() was run with the dataset-specific variable genes, peak_to_gene_method = “Signac”, and parallel = TRUE. Unspecified arguments used Pando v1.1.1 defaults, including Gaussian generalized linear models, candidate regions overlapping each annotated gene body or its strand-aware 100-kb upstream extension, an absolute TF-target expression-correlation threshold of 0.1, a peak-target accessibility-correlation threshold of 0, unscaled model inputs, and FDR adjustment of coefficient *p* values. Each modeled coefficient represented an interaction between accessibility of a motif-containing candidate region and expression of the corresponding motif-matched TF in relation to target-gene expression.

During inference, OpenMP and BLAS threads were restricted to one before registering a four-worker PSOCK cluster for Pando, to avoid CPU oversubscription while retaining independent worker processes. After inference, complete coefficient and model-fit tables were retained. Region-level associations with FDR-adjusted *p* < 0.05 were considered supported and were classified as positive or negative according to the sign of the coefficient estimate. Pando modules were identified using find_modules(p_thresh = 0.05, rsq_thresh = 0.05, nvar_thresh = 2, min_genes_per_module = 1).

#### Cross-cell-type GRN graph and union/intersection analysis

Cross-dataset graph analysis was performed after independent Pando model fitting for CTL, HDF, and HAEC and did not involve refitting a pooled model. To construct TF-to-TF backbone networks, supported positive associations were restricted to edges for which both the source TF and target gene belonged to the Pando TF universe. Repeated region-level support for the same directed TF-target pair within each dataset was collapsed to one directed edge, retaining the maximum and mean coefficient estimates and the number of supporting region-level records.

Directed graph objects were constructed using tidygraph/igraph and visualized with ggraph. The union of the three cell-type-specific TF-to-TF networks was used as a shared reference edge universe and contained every unique directed TF-to-TF relationship detected in at least one of CTL, HDF, or HAEC. Each union edge was annotated by its exact three-cell-type presence pattern, with directionality preserved such that TF-A-to-TF-B and TF-B-to-TF-A were treated as distinct edges. This representation was used to extract the strict three-cell-type intersection, relaxed intersections supported by at least two cell types, and pairwise intersections.

In-degree, out-degree, total degree, and coefficient summaries were recalculated within the shared graph representations to prioritize TFs with recurrent incoming or outgoing connectivity across cell types. Shared edges were overlaid on the complete union topology to distinguish recurrent TF hubs and cross-cell-type core regulatory structures from cell-type-restricted branches. Accordingly, TFs nominated by this analysis were prioritized based on recurrent cross-cell-type network connectivity rather than high expression or connectivity in a single cell type alone, and these graph-based nominations were not interpreted as direct causal validation.

#### Pando TF-regulatory region-target gene analysis

We next extended the cross-cell-type comparison from TF-to-TF relationships to all supported positive TF-to-target relationships, including non-TF target genes. Because multiple regulatory regions could support the same TF-target relationship, region-level records were collapsed to one directed TF-to-target edge for gene-level comparison. A shared gene-level edge required the same regulator TF, the same target gene, and the same edge direction in multiple cell types. Each edge in the union network was classified as detected in one, two, or all three cell types, from which pairwise and three-cell-type intersections were obtained.

We then asked whether shared TF-to-target relationships were supported by the same genomic regulatory regions across cell types. Because overlapping Pando regions could have slightly different coordinates, regions from all three cell types were merged into a common set of non-overlapping consensus intervals using GenomicRanges::reduce(). Each original Pando region was assigned to the consensus interval that it overlapped. A regulatory-region-level association was considered shared only when the same TF, the same target gene, and the same consensus interval were detected in multiple cell types. This criterion was more stringent than gene-level sharing, which required only the same directed TF-to-target relationship.

Results were visualized as TF-to-target graphs and as three-layer TF-consensus-region-target graphs. The two levels of sharing were therefore reported separately: a TF-to-target relationship detected in all three cell types was classified as gene-level shared, whereas it was classified as regulatory-region-level shared only if at least one matching consensus interval also supported that relationship across the same cell types. Consensus intervals were computationally defined and were not interpreted as experimentally validated enhancers.

#### TOX-centered target and CRM analysis

For *TOX*-focused analysis of the Pando GRNs, positive *TOX* associations (FDR-adjusted *p* < 0.05 and estimate > 0) were further evaluated for cluster-level RNA enrichment with Seurat FindMarkers() using dataset-specific Leiden cluster assignments. Each selected Leiden cluster was compared with the remaining selected clusters, and the tested features were restricted to positive Pando-supported *TOX* targets. Thus, FindMarkers() was not used to discover *TOX* targets de novo from the transcriptome, but to determine whether Pando-supported targets showed higher RNA expression in a selected cluster relative to the remaining selected clusters. The default Wilcoxon rank-sum test was run with only.pos = TRUE, min.pct = 0.05, and logfc.threshold = 0.1. The cluster-level output retained the Leiden cluster identity and RNA fold-change information and was joined to the Pando-supported target list. Genes retained by this expression filter in at least one tested cluster were termed cluster-supported *TOX* candidate targets. A gene could occur in multiple rows when enriched in multiple clusters; these records were collapsed by gene identity before cross-dataset comparison. Sharing classes were then defined by whether each unique gene occurred in one, two, or all three candidate tables and were calculated separately from sharing of the underlying CRMs.

The *TOX*-centered tripartite network was generated by restricting positive Pando TF-region-target associations to the cluster-supported *TOX* candidate genes and then mapping the regulatory regions to consensus CRMs. *TOX* was displayed as the focal TF. Other TFs were included as potential co-regulators when a positive Pando TF-region-target association involving the same candidate gene was mapped to a consensus CRM. The three layers represented TFs, computationally defined consensus CRMs, and candidate target genes. Each consensus CRM was generated by merging overlapping Pando regulatory regions across the three datasets and served as a common genomic unit for comparing TF-CRM-target connections across cell types. Pando mapped three JASPAR motifs to *TOX* by sequence similarity (MA0077.1, MA0442.2, and MA0523.1). A consensus CRM was considered *TOX*-associated when at least one underlying Pando region contained one of these *TOX*-mapped motifs and the corresponding *TOX*-region-target coefficient was positive with FDR-adjusted *p* < 0.05. Target-node classes were determined solely by whether a gene occurred in one, two, or all three of the CTL, HDF, and HAEC candidate tables. For each TF-consensus-CRM-target connection, the edge table recorded the number of cell types supporting the same TF, target gene, and consensus CRM. In the final visualization, all CRM nodes and edges were displayed uniformly in gray; target-node color therefore represented candidate-gene sharing rather than CRM sharing.

To identify KD-supported TOX downstream candidate genes for Figure 7B, the cluster-supported TOX candidate targets defined above were intersected with the corresponding cell type-matched bulk RNA-seq genes that decreased after *TOX* knockdown. KD support was defined by a negative log2 fold change after *TOX* depletion relative to the matched control, without applying an additional FDR threshold. The resulting gene lists therefore required Pando-supported TOX regulatory association, cluster-level RNA enrichment, and concordant reduction after *TOX* depletion. Unique genes passing this combined filter were used for the HDF, CTL, and HAEC Venn analysis in Figure 7B and downstream functional summaries. Sharing classes were defined by whether each unique KD-supported candidate gene was present in one, two, or all three cell-type-specific filtered lists.

#### scPrinter/seq2PRINT footprint and sequence-attribution analysis

Single-cell chromatin accessibility profiles from HDF, CTL, and HAEC were analyzed using hg38-based scPrinter/seq2PRINT models^36^. Processed scPrinter objects contained cell metadata, insertion profiles, aggregate accessibility tracks, and peak definitions. Cell-state or stage labels used for stage-specific low-rank adaptation (LoRA) fine-tuning were inherited from the preceding ArchR-associated annotation workflow and stored in the scPrinter metadata. HDF stages were defined as d0, d7c, d7g1, d7g2, and d7g3; CTL stages as d0, d5c, d5g1, d5g2, and d5g3; and HAEC stages as d0, d7c, d7g1, and d7g2. Fragment files were matched to annotated cells during scPrinter object construction. Candidate regulatory regions were generated by all-cell scPrinter/MACS2 peak calling and converted into seq2PRINT-compatible BED files.

For each dataset, bulk scPrinter models were first trained to predict aggregate chromatin-accessibility profiles from DNA sequence using a five-fold chromosome-held-out split. Training used reverse-complement augmentation, positional jittering, convolutional and inception blocks, batch normalization, exponential moving average weights, and mixed-precision training. Fold-specific bulk checkpoints were then used to initialize the corresponding stage-specific LoRA fine-tuning models. Bulk training and LoRA fine-tuning were performed on a 48 GB NVIDIA RTX 6000 Ada GPU.

Stage-resolved sequence attribution was computed for candidate *TOX* and *COL25A1* regulatory loci using the fine-tuned LoRA models. Candidate peaks and visualization windows were selected from peak-to-gene links assigned to the corresponding target genes and then prioritized for focused visualization based on concordant stage-associated attribution dynamics across HDF, CTL, and HAEC. For *TOX*, the full gene locus together with flanking intergenic regions was inspected, and seven regulatory windows with reproducible stage-associated attribution patterns were retained for visualization. Hypothetical sequence-attribution scores were calculated for each fold and stage, fold-level profiles were averaged, and dynamic seqlet regions were visualized as sequence-logo tracks. Motif annotations for *TOX* plots were obtained from the UCSC-hosted JASPAR 2026 track, with labels prioritized for hits overlapping highlighted seqlet regions and redundant labels suppressed.

For *COL25A1*, two representative peak-to-gene-linked peaks with concordant attribution dynamics across HDF, CTL, and HAEC were selected for visualization. LoRA attribution profiles were plotted for these regions, and motif matches were annotated using a custom FIMO scan restricted to the three Pando-assigned TOX-associated motif classes: HMG/SOX9-like, HMG/SOX10-like, and TCF/TCF7L2-like. Final highlight windows and motif labels were assigned by integrating exact FIMO hit positions with dynamic scPrinter seqlet patterns.

#### Public scRNA-seq dataset reanalysis

Publicly available single-cell RNA-seq datasets of human fibroblast reprogramming were downloaded from GEO, including SeV-OSKM (GSE242424; scRNA-seq subseries GSE242423), lentiviral OSKM (GSE118258), and mRNA OSKMLN (GSE221739) datasets. When processed count matrices were available, they were used directly; otherwise, raw matrix files were assembled into gene-by-cell count matrices. Downstream analysis was performed in R using Seurat. Cells were normalized, variable features were identified, and principal component analysis was performed, followed by nearest-neighbor graph construction and UMAP visualization. Cells were annotated by the time point information provided in the original datasets. *TOX* expression was visualized on UMAP embeddings using normalized expression values, and dot plots were generated to summarize average *TOX* expression and the percentage of *TOX*-expressing cells at each time point.

#### *TOX* loss-and gain-of-function during reprogramming

For siRNA-mediated *TOX* knockdown, HDFs and HAECs were transfected with 20 nM of one of three *TOX*-targeting siRNAs or a medium-GC non-targeting negative-control siRNA (siCtrl) using Lipofectamine RNAiMAX in Opti-MEM on days 1 and 4 after SeV-OSKM transduction. The transfection medium was replaced with the corresponding cell type-specific medium after 5 h. Cells were replated on day 5, and RNA was collected on day 7. For iPSC colony-formation assays, siRNA transfection was repeated on days 7 and 10, and colonies were counted on day 14.

For shRNA-mediated *TOX* knockdown, CTLs were transduced with one of three lentiviral vectors expressing *TOX*-targeting shRNAs or a lentiviral vector expressing a scrambled-sequence control shRNA (shCtrl) and selected with 1 µg/mL puromycin for 7 days. Following selection, cells were stimulated with CD3/CD28 Dynabeads on day −4, transduced with SeV-OSKM on day 0, and replated on day 1. RNA was collected on day 5, and iPSC colonies were counted on day 14.

siTOX #1 was used for bulk RNA-seq analyses in HDFs and HAECs, and shTOX #1 was used in CTLs. For reprogramming-independent controls, HDFs and HAECs were treated with the same *TOX*-targeting siRNAs or siCtrl in the absence of SeV-OSKM; *TOX* mRNA and total recovered cell numbers were measured 2 and 3 days after transfection, respectively. CTLs transduced with the same shTOX or shCtrl lentiviral vectors were selected with 1 µg/mL puromycin for 7 days and then stimulated with CD3/CD28 Dynabeads without SeV-OSKM transduction; *TOX* mRNA and total recovered cell numbers were assessed 3 days after stimulation.

At the day 7 RNA-collection time point for HDFs and HAECs and the day 5 time point for CTLs, the absolute yield of each Gr population was estimated by multiplying its flow-defined fraction among recovered parent cells by the total recovered cell count. For the relative-yield analysis, donor-matched log2(siTOX/siCtrl) values for HDFs and HAECs or log2(shTOX/shCtrl) values for CTLs were calculated separately for each Gr. Differences among relative Gr-cell yields were assessed by two-way repeated-measures ANOVA with matching by donor across both Gr state and siRNA/shRNA identity and with Geisser-Greenhouse correction. Gr2 and Gr3 were compared with Gr1 within each reagent using Dunnett-adjusted comparisons for HDFs and CTLs; Gr2 was compared with Gr1 within each siRNA using Sidak-adjusted comparisons for HAECs.

For *TOX* gain-of-function experiments, HDFs were co-transduced with a doxycycline-inducible human *TOX* lentiviral vector or the corresponding ORF_Stuffer control vector and a tTS/rtTA lentiviral vector. Cells were selected with 1 µg/mL puromycin and 6 µg/mL blasticidin for 7 days before SeV-OSKM transduction. *TOX* expression was induced with 0.1 µg/mL doxycycline during the indicated time windows. For transcriptomic analysis, doxycycline was administered during days 1–7 or days 4–7, and RNA was collected on day 7. For iPSC colony-formation assays, doxycycline was administered during days 1–14, 1–10, 4–10, 7–10, or 10–14; a doxycycline-free condition was included as a control. Cells were replated on day 5, and colonies were counted on day 14.

### QUANTIFICATION AND STATISTICAL ANALYSIS

Statistical analyses were performed using GraphPad Prism 10 unless otherwise indicated. Data from four independent donors per cell type are shown as mean ± SD together with donor-level values. Two-condition matched comparisons used two-tailed paired t tests. Comparisons across more than two matched conditions used repeated-measures one-way ANOVA followed by Dunnett’s multiple-comparisons test versus the indicated control. For visualization, bulk RNA-seq module scores were expressed as within-donor differences from matched Ctrl values, such that Ctrl was zero by definition.

Statistical comparisons and the significance values shown in the figures were calculated from the corresponding original (uncentered) donor-level scores. Differential expression, multiple-testing correction, and gene set enrichment analyses were performed as described above. The statistical test and significance notation used for each experiment are specified in the corresponding figure legend.

## ACKNOWLEDGMENTS

We are grateful to M. Yamanaka, Y. Tsukamoto, H. Kunitomi, and K. Tomoda for sharing data and for helpful discussions prior to publication; Kevin Pastores and Jane Srivastava for assistance with cell sorting; Ayushi Agrawal and Reuben Thomas for assistance with DNA methylation array data analysis; Yan Hu for guidance on scPrinter-based analyses; Jitsutaro Kawaguchi for providing the customized SeV vectors; and Françoise Chanut for insightful feedback on the framing, organization, and presentation of the manuscript, as well as for editorial assistance and proofreading. This work was supported by Altos Labs and the California Institute for Regenerative Medicine (CIRM). Additional support was provided by Mr. Hiroshi Mikitani, Mr. Marc Benioff, the L. K. Whittier Foundation, the Roddenberry Foundation, Gladstone Institutes, and the National Heart, Lung, and Blood Institute of the National Institutes of Health (U01-HL100406 and U01-HL098179).

## AUTHOR CONTRIBUTIONS

K.W. performed most of the HDF experiments and analyzed the resulting data. X.W. performed most of the single-nucleus multiome analyses. J.M.D. performed most of the CTL experiments. S.Y. supervised the study. A.K. conceived and designed the study, performed most of the experimental work, with a particular focus on HAECs, analyzed data, supervised the research, and wrote the manuscript.

---

**Figure S1.**
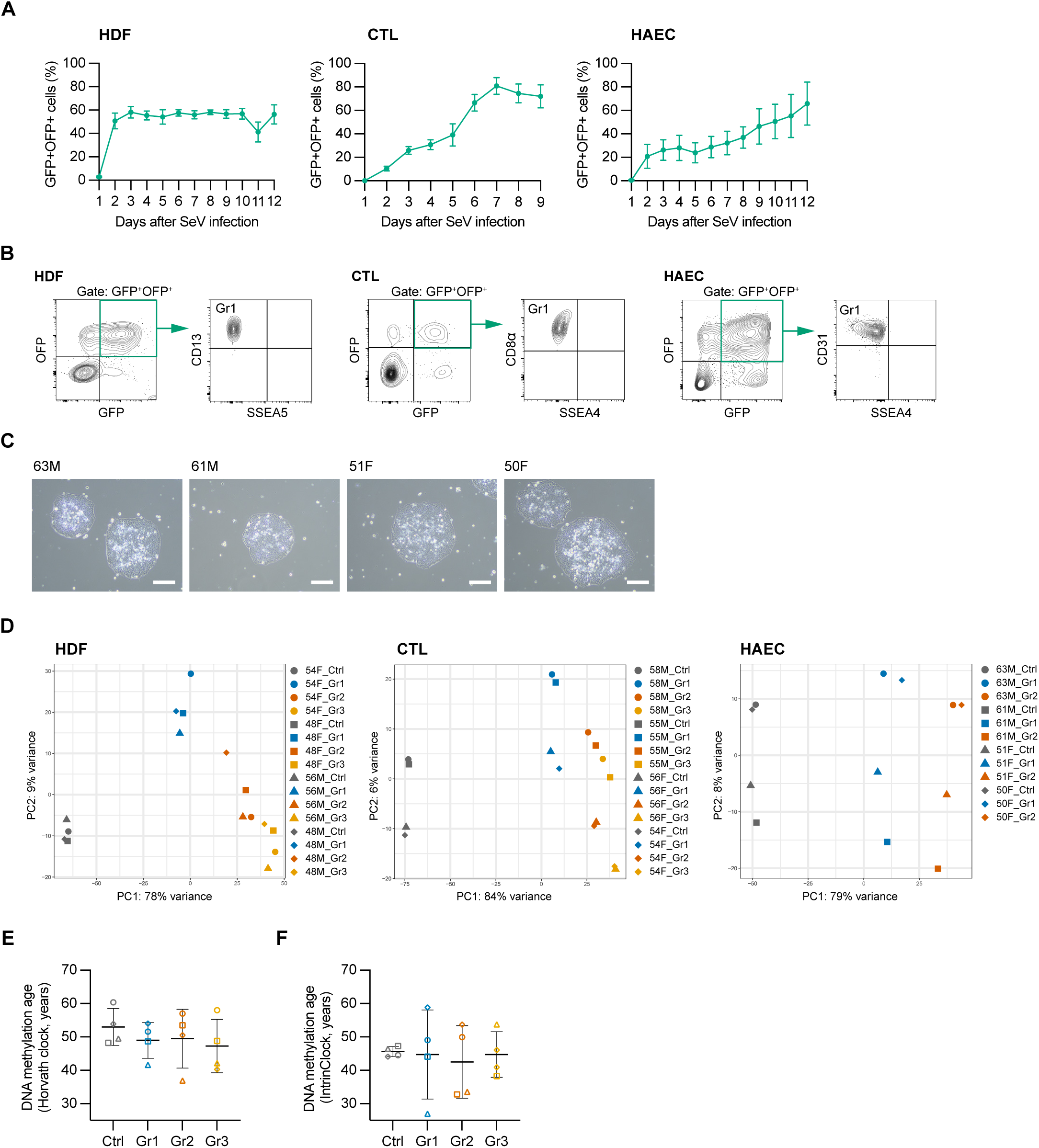
Supporting validation of the fluorescent temperature-sensitive SeV-OSKM system and early molecular profiling. (A) Time-course quantification of GFP+OFP+ cells after SeV-OSKM infection in HDFs, CTLs, and HAECs. Line plots show mean values from four independent donors with SD. (B) Representative flow cytometry plots of matched Ctrl-vector-infected HDFs, CTLs, and HAECs at the sorting time point. GFP+OFP+ cells were evaluated using CD13 and SSEA5 for HDFs, CD8α and SSEA4 for CTLs, and CD31 and SSEA4 for HAECs. In each cell type, the Ctrl population retained the Gr1 marker profile. (C) Representative iPSC colonies derived from HAEC donors. Scale bars, 200 µm. (D) PCA of bulk RNA-seq profiles from sorted groups in each cell type. Samples separate according to reprogramming state in each cell type. (E) DNA methylation age estimates in CTLs using the Horvath clock, performed as a complementary analysis to the BLUP clock analysis shown in Figure 1H. The central horizontal line indicates the mean of four independent donors, and error bars indicate SD; open symbols indicate individual CTL donors: circle, 58M; square, 55M; triangle, 56F; and diamond, 54F. Repeated-measures one-way ANOVA followed by Dunnett’s multiple comparisons test versus Ctrl was used where indicated; unmarked comparisons are not significant. (F) DNA methylation age estimates in CTLs using IntrinClock, performed as a complementary analysis to the BLUP clock analysis shown in Figure 1H. Plotting and statistical conventions are as in panel E.

**Figure S2.**
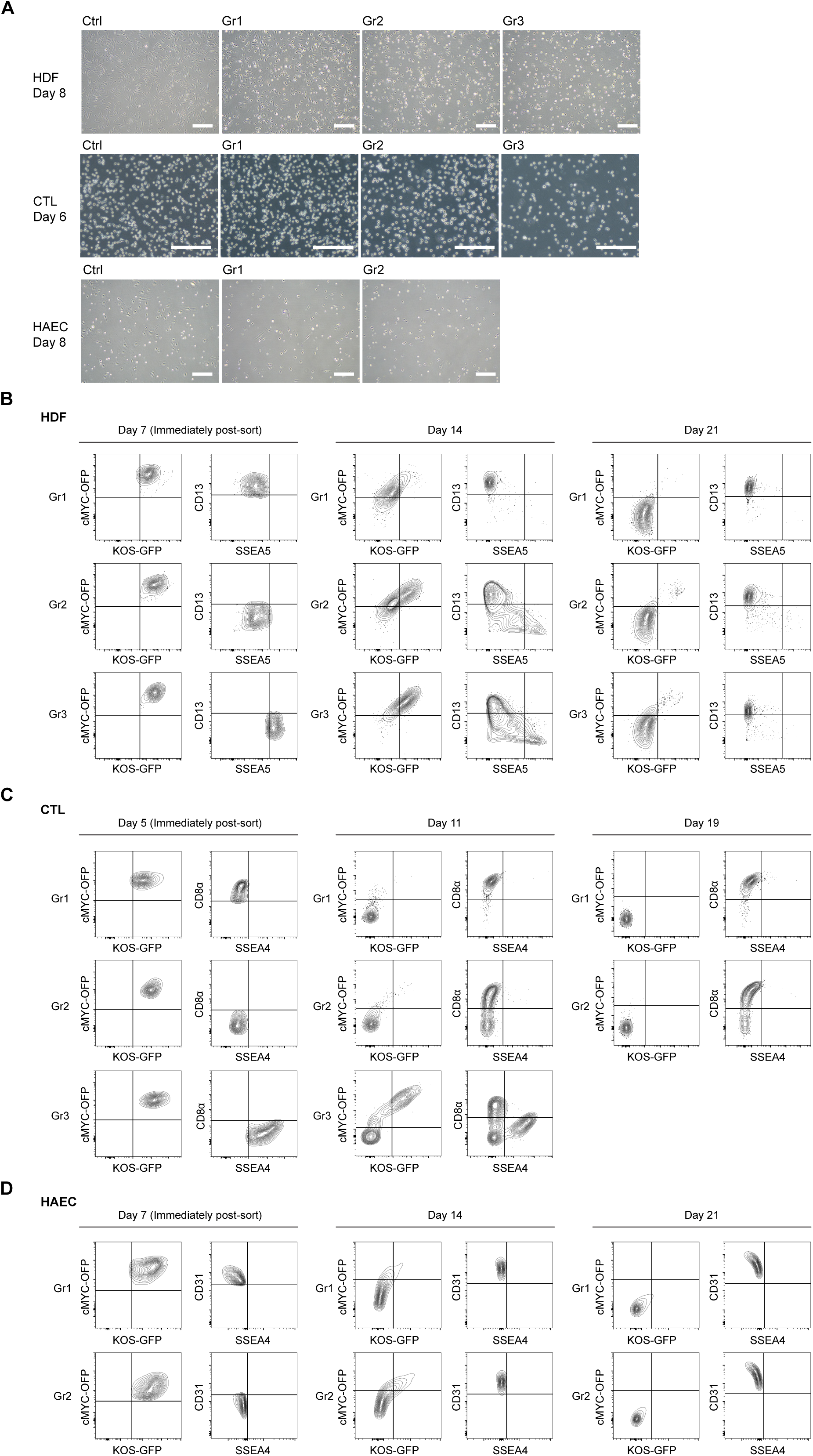
Early morphology and marker recovery after sorting. (A) Representative microscopy images of HDF, CTL, and HAEC groups on the day after sorting and temperature shift. HDF and HAEC groups show morphological changes corresponding to reprogramming depth, whereas CTL morphology is less distinct and CTL Gr3 shows reduced cell number. Scale bars, 200 µm. (B) Representative flow cytometry plots showing longitudinal changes in KOS-GFP, cMYC-OFP, SSEA5, and CD13 in HDF groups after sorting. These plots complement the quantitative recovery analysis shown in Figure 2D. (C) Representative flow cytometry plots showing longitudinal changes in KOS-GFP, cMYC-OFP, SSEA4, and CD8α in CTL groups after sorting. These plots complement the quantitative recovery analysis shown in Figure 2F. (D) Representative flow cytometry plots showing longitudinal changes in KOS-GFP, cMYC-OFP, SSEA4, and CD31 in HAEC Gr1 and Gr2 groups after sorting. These plots complement the quantitative recovery analysis shown in Figure 2H. HAEC Gr3 was not analyzed because of insufficient cell numbers.

**Figure S3.**
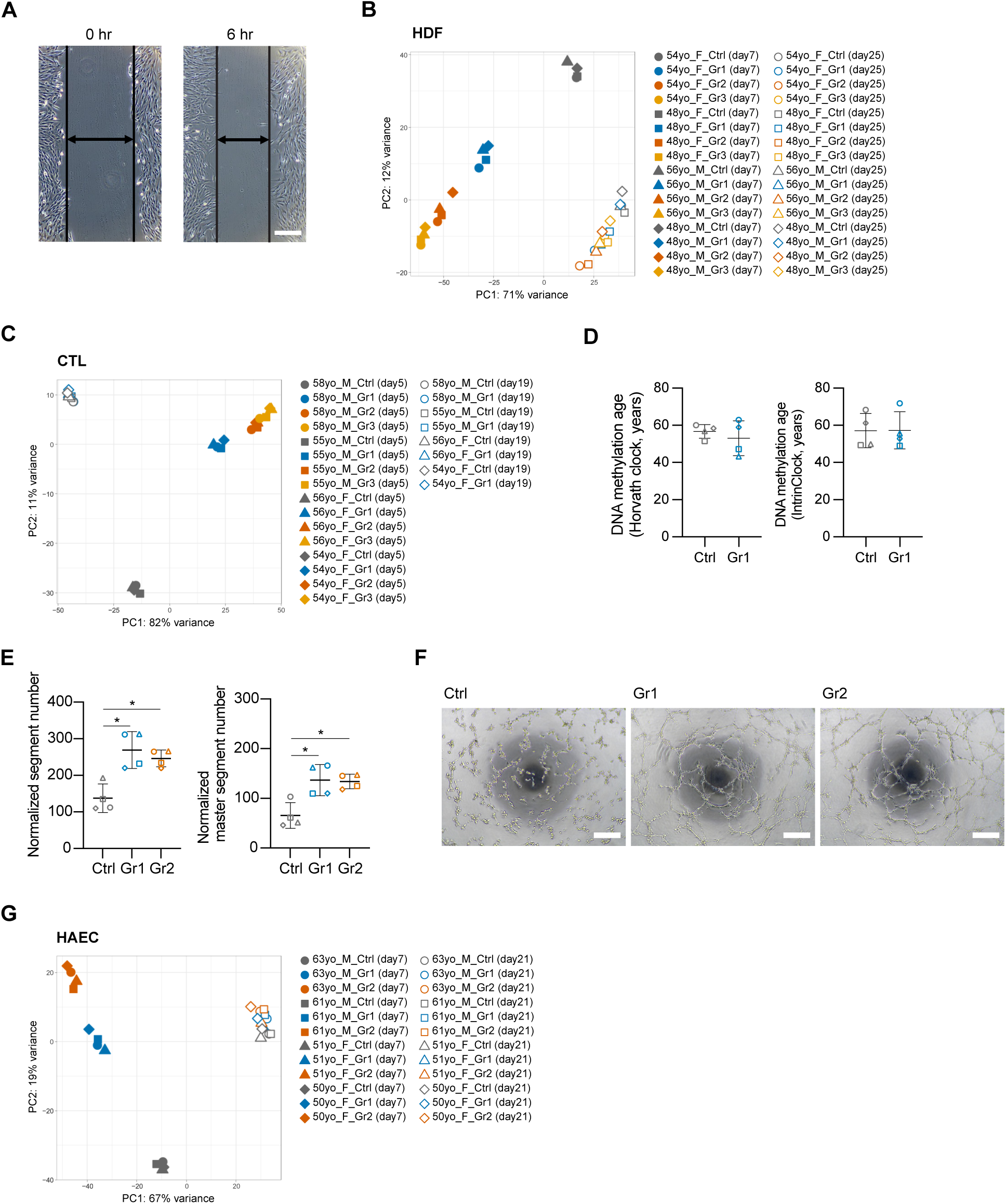
Supporting assays for endpoint recovery and function after partial reprogramming. (A) Representative scratch wound-healing images in HDFs at 0 hr and 6 hr. Scale bars, 200 µm. Quantification is shown in Figure 3C. (B) PCA of HDF bulk RNA-seq profiles before and after temperature-shift-mediated suppression of SeV-OSKM expression. Day 7 samples separated strongly by reprogramming state, whereas day 25 samples converged toward a shared somatic transcriptomic state. (C) PCA of CTL bulk RNA-seq profiles before and after temperature-shift-mediated suppression of SeV-OSKM expression. Day 19 Ctrl and Gr1 samples show convergence relative to the day 5 sorted states. (D) DNA methylation age estimates in CTLs at day 26 using the Horvath clock and IntrinClock, in addition to the BLUP clock analysis shown in Figure 3H. No significant difference between Ctrl and Gr1 was detected by either clock. The central horizontal line indicates the mean of four independent donors, and error bars indicate SD; open symbols indicate CTL donors as in Figure 3F. A two-tailed paired t test was used; unmarked comparisons are not significant. (E) Cell-number-normalized segment number and master segment number in HAEC tube-formation assays. The central horizontal line indicates the mean of four independent donors, and error bars indicate SD; open symbols indicate the HAEC donors listed in Figure 3K. Repeated-measures one-way ANOVA followed by Dunnett’s multiple comparisons test versus Ctrl was used. A prespecified comparison between Gr1 and Gr2 was additionally performed using a Sidak-adjusted comparison; no significant difference was detected. \**p* < 0.05; unmarked comparisons are not significant. (F) Representative tube formation images for HAEC Ctrl, Gr1, and Gr2. Gr1 and Gr2 showed more extensive tube-like network formation than Ctrl. Scale bars, 200 µm. Quantification is shown in Figure 3L and Figure S3E. (G) PCA of HAEC bulk RNA-seq profiles before and after temperature-shift-mediated suppression of SeV-OSKM expression. Day 21 samples show convergence relative to the day 7 sorted states.

**Figure S4.**
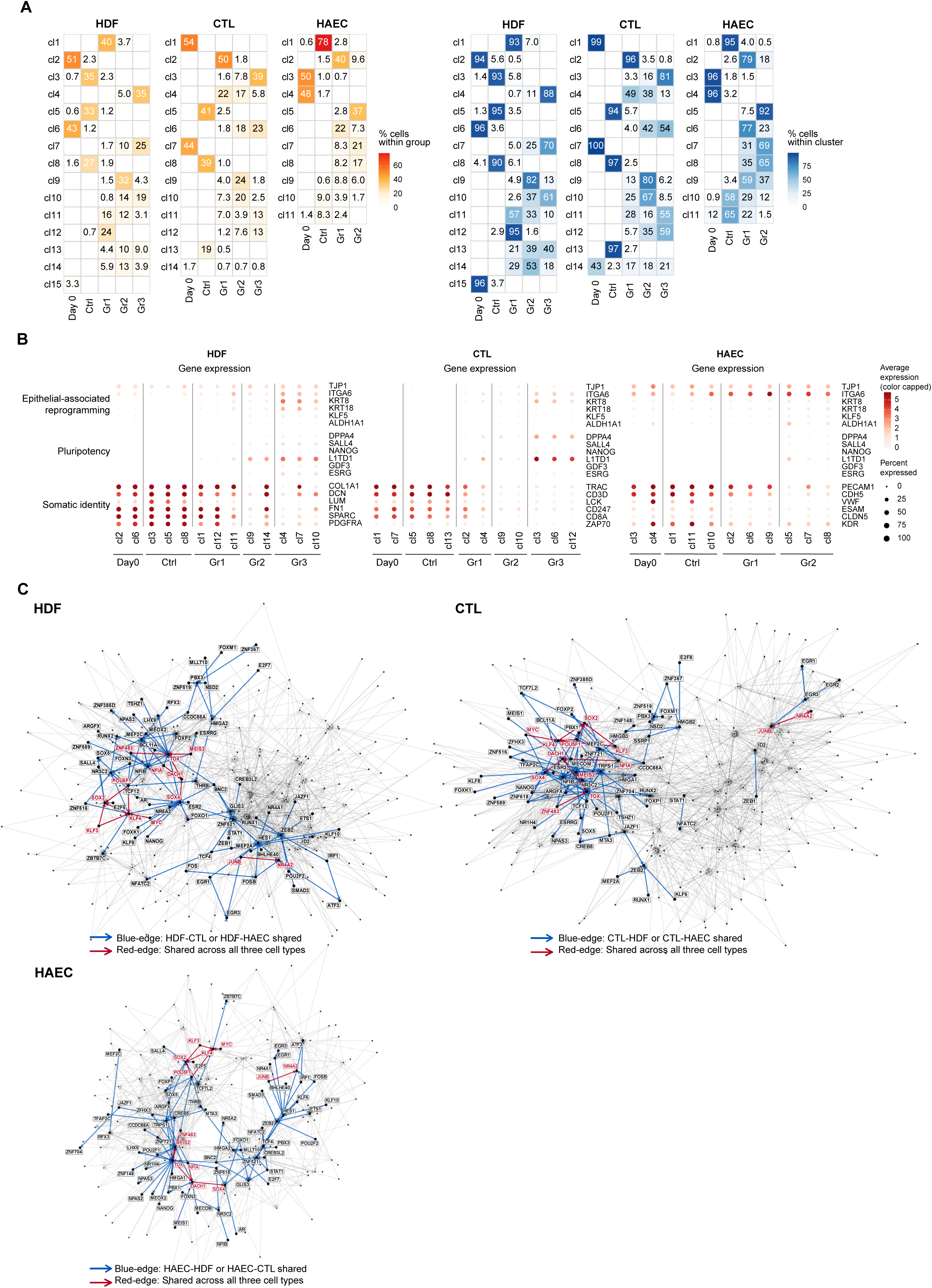
Supporting single-nucleus multiome analyses. (A) Fraction of cells from each flow-sorted group within representative Leiden clusters and fraction of cells within each cluster contributed by each group. These summaries were used to select representative clusters for downstream linked peak and gene expression analyses. (B) Dot plots of representative epithelial-associated reprogramming (MET-associated), pluripotency, and somatic identity gene expression across clusters in HDFs, CTLs, and HAECs. Dot size indicates the fraction of cells expressing each gene and color indicates average expression with capped color scaling. (C) Pando-inferred transcription factor network modules shared between pairs of cell types and across all three cell types. Blue edges denote regulatory edges shared by exactly two cell types, and red edges denote regulatory edges shared across all three cell types. TOX appears among shared reprogramming-associated regulatory modules.

**Figure S5.**
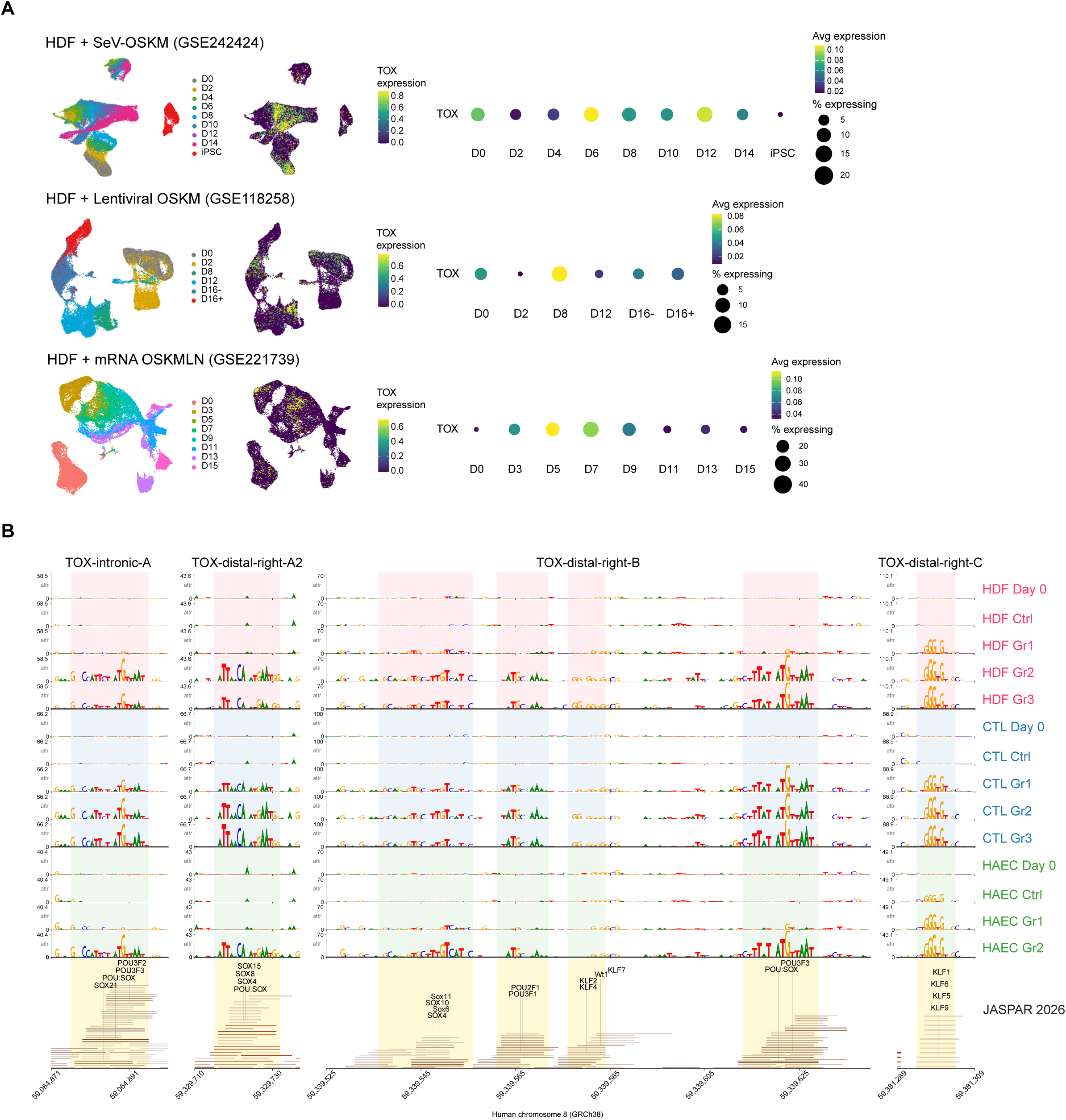
Transient *TOX* induction in public human fibroblast reprogramming datasets and additional *TOX*-linked candidate cis-regulatory elements. (A) Reanalysis of three public human fibroblast reprogramming single-cell RNA-seq datasets generated using SeV-OSKM, lentiviral OSKM, or mRNA OSKMLN. UMAPs show reprogramming time points and *TOX* RNA expression, and dot plots summarize *TOX* expression over time. Dot color indicates average *TOX* expression and dot size indicates the percentage of *TOX*-expressing cells. *TOX* was transiently induced during early-to-mid reprogramming across all three platforms. (B) State-resolved scPrinter/seq2PRINT sequence-attribution profiles and JASPAR 2026 motif annotations for the four remaining labeled *TOX*-linked candidate CREs: *TOX*-intronic-A, *TOX*-distal-right-A2, *TOX*-distal-right-B, and *TOX*-distal-right-C. Together with the three elements shown in Figure 5E, this panel displays all seven candidate CREs highlighted in Figure 5D.

**Figure S6.**
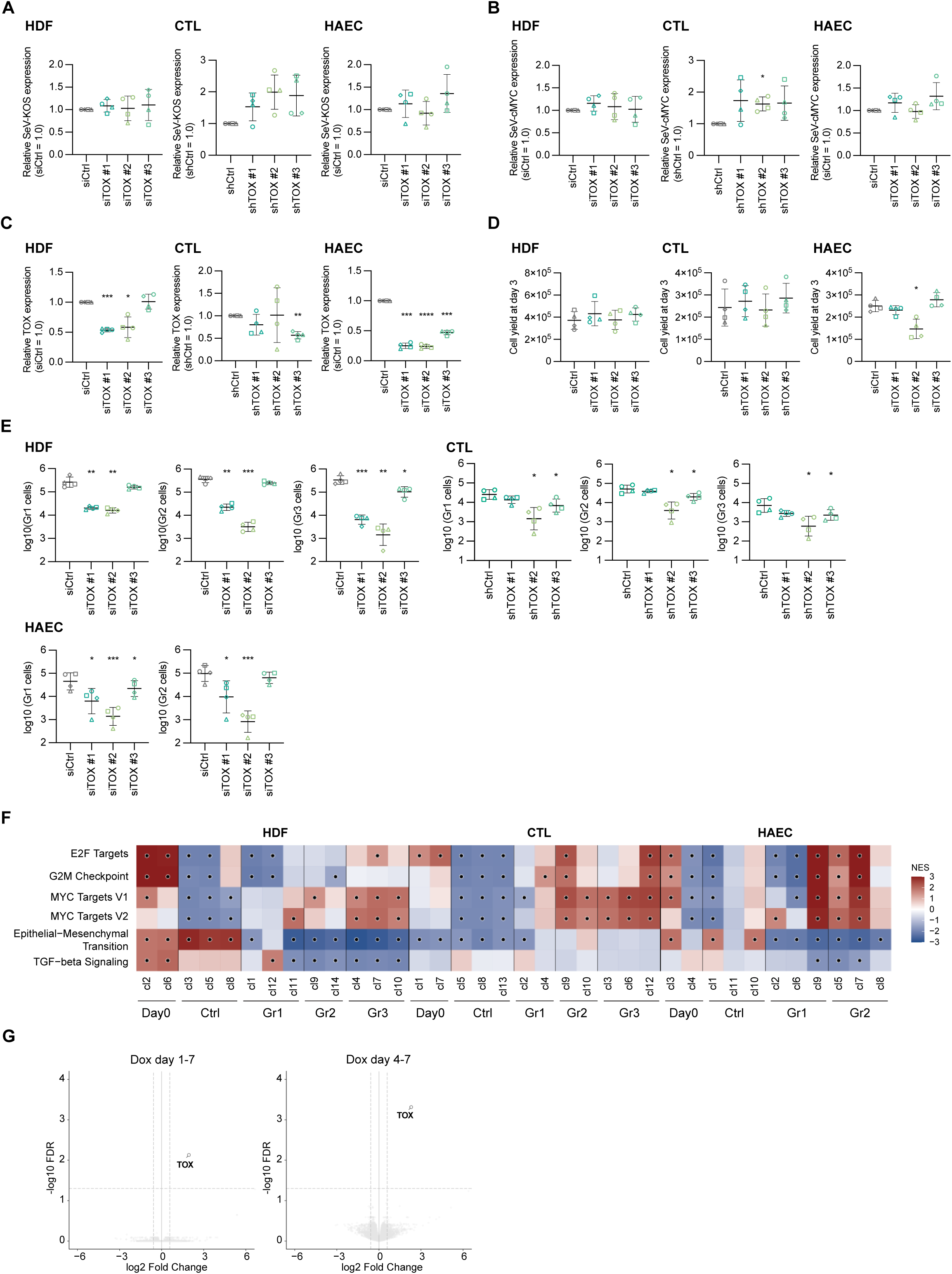
Supporting controls, absolute Gr-cell yields, state-resolved pathway analyses, and *TOX*-overexpression transcriptomic results. (A) SeV-KOS expression measured by qPCR on day 2 after SeV-OSKM transduction and *TOX* knockdown. The central horizontal line indicates the mean of four independent donors, and error bars indicate SD; open symbols indicate the donors listed in Figure 6B. Repeated-measures one-way ANOVA followed by Dunnett’s multiple comparisons test versus the matched control (siCtrl for HDFs and HAECs; shCtrl for CTLs) was used where indicated. Unmarked comparisons are not significant. (B) SeV-cMYC expression measured by qPCR on day 2 after SeV-OSKM transduction and *TOX* knockdown. The central horizontal line indicates the mean of four independent donors, and error bars indicate SD; open symbols indicate the donors listed in Figure 6B. Repeated-measures one-way ANOVA followed by Dunnett’s multiple comparisons test versus the matched control (siCtrl for HDFs and HAECs; shCtrl for CTLs) was used where indicated. \**p* < 0.05; unmarked comparisons are not significant. (C) Relative *TOX* expression measured by qPCR under control conditions without SeV-OSKM. For HDFs and HAECs, RNA was collected 2 days after transfection with one of the *TOX*-targeting siRNAs or siCtrl. CTLs transduced with a shTOX or shCtrl lentiviral vector were selected with puromycin for 7 days and then stimulated with CD3/CD28 Dynabeads; RNA was collected 3 days after stimulation. *TOX* expression in CTLs remained very low under this condition, resulting in variable relative estimates. The central horizontal line indicates the mean of four independent donors, and error bars indicate SD; open symbols indicate individual donors: HDFs, circle 54F, square 48F, triangle 56M, diamond 48M; CTLs, circle 58M, square 55M, triangle 56F, diamond 54F; HAECs, circle 63M, square 61M, triangle 51F, diamond 50F. Repeated-measures one-way ANOVA followed by Dunnett’s multiple comparisons test versus siCtrl for HDFs and HAECs or shCtrl for CTLs was used. \**p* < 0.05, \*\**p* < 0.01, \*\*\**p* < 0.001, \*\*\*\**p* < 0.0001; unmarked comparisons are not significant. (D) Total cell recovery under the corresponding control conditions without SeV-OSKM. HDFs and HAECs were counted 3 days after siRNA transfection. Puromycin-selected CTLs transduced with a shTOX or shCtrl lentiviral vector were counted 3 days after CD3/CD28 stimulation. The central horizontal line indicates the mean of four independent donors, and error bars indicate SD; open symbols indicate the donors listed in Figure 6B. Repeated-measures one-way ANOVA followed by Dunnett’s multiple comparisons test versus siCtrl for HDFs and HAECs or shCtrl for CTLs was used. \**p* < 0.05; unmarked comparisons are not significant. (E) Absolute yields of marker-defined Gr populations after *TOX* knockdown at the day 7 analysis time point for HDFs and HAECs and day 5 for CTLs. Values are shown as log10 cell number for each Gr. The central horizontal line indicates the mean of four independent donors, and error bars indicate SD; open symbols indicate the donors listed in Figure 6B. Within each Gr, repeated-measures one-way ANOVA followed by Dunnett’s multiple-comparisons test versus the corresponding siCtrl or shCtrl was used. \**p* < 0.05, \*\**p* < 0.01, \*\*\**p* < 0.001; unmarked comparisons are not significant. (F) GSEA heatmap across representative single-nucleus multiome clusters spanning the graded reprogramming states, showing pathway activity during unperturbed reprogramming progression. This analysis provides a state-resolved reference for interpreting the pathway shifts observed after *TOX* knockdown in Figure 6G. Black dots indicate significantly enriched gene sets with FDR < 0.05. (G) Volcano plots of bulk RNA-seq after doxycycline-inducible *TOX* overexpression during days 1-7 or days 4-7 of reprogramming. Genes with FDR < 0.05 and |log2 fold change| > 1 were considered significantly differentially expressed; dashed lines indicate these thresholds. *TOX* was the dominant upregulated gene at the individual-gene level; pathway-level results are shown in Figure 6K.

**Figure S7.**
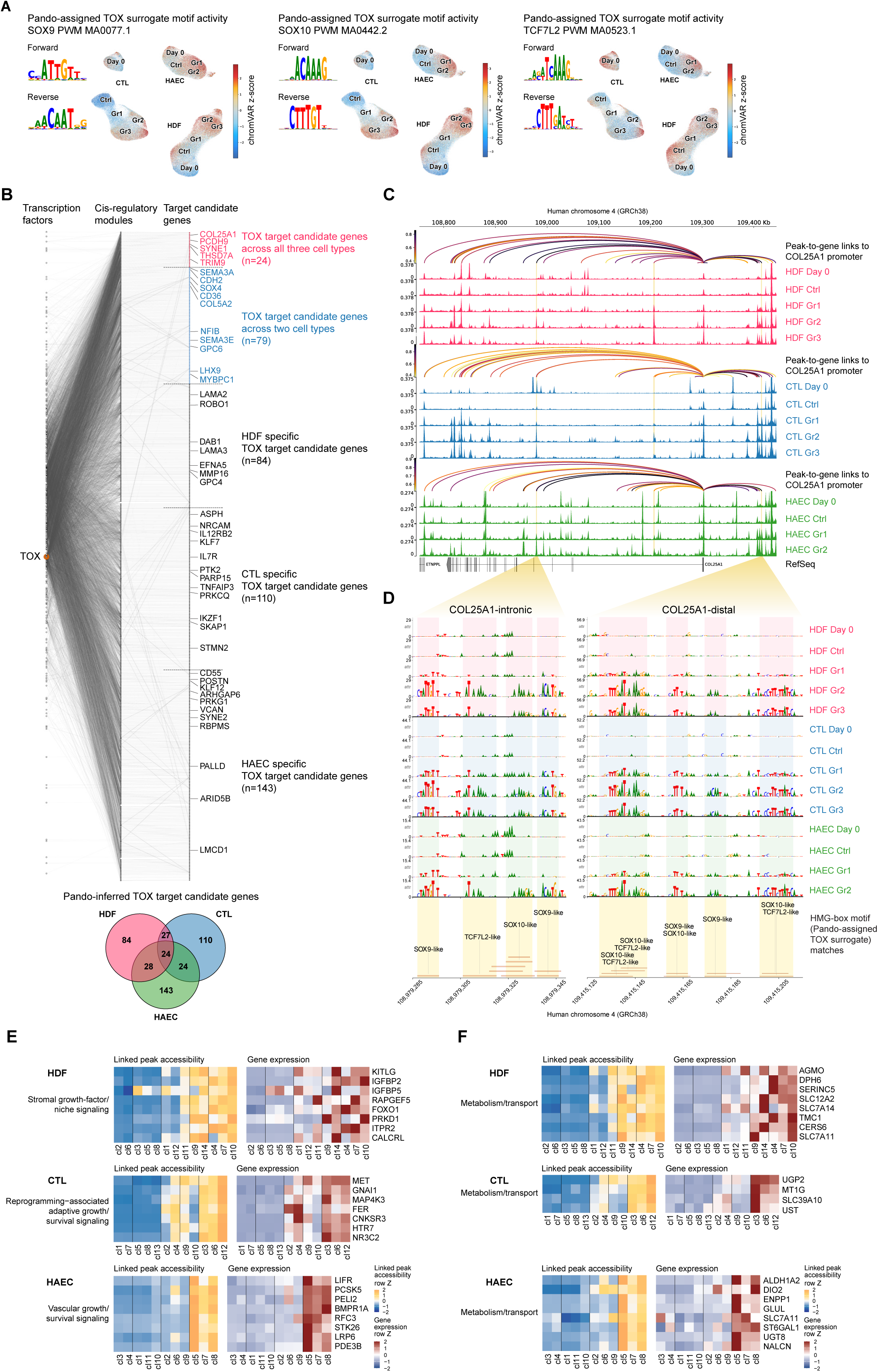
Pando-assigned TOX surrogate motifs, network structure, and supporting analyses of Pando-inferred TOX-associated candidate downstream programs. (A) Pando-assigned TOX surrogate motifs. Pando’s motif reference contains no TOX-specific position weight matrix; instead, its default motif-to-TF map assigns TOX three PWMs of related HMG-box factors on the basis of protein-sequence similarity (SOX9 MA0077.1, SOX10 MA0442.2, and TCF7L2 MA0523.1), and these assignments were used without modification. Forward and reverse-complement sequence logos are shown for each PWM, together with chromVAR deviation Z-scores for that motif projected onto the integrated ATAC UMAP across HDFs, CTLs, and HAECs. The three motifs gave broadly related but model-dependent patterns of activity across reprogramming states. In each panel the color scale spans the 10th to 90th percentile of the chromVAR Z-score distribution for that motif, and values outside this range are shown at the limiting colors. Because these scores are not TOX-specific and may also reflect accessibility associated with other HMG-box factors, including exogenous SOX2, they were interpreted as motif-associated chromatin accessibility rather than direct TOX occupancy, and candidate target genes were instead filtered by *TOX* knockdown (Figure 7A). (B) Pando network overview and pre-filter overlap of inferred TOX target candidate genes. The upper network shows inferred transcription factor-CRE module-target gene relationships. TOX is highlighted, and target genes included in Figures 7E and 7F are labeled and grouped as shared across all three cell types, shared across exactly two cell types, or cell type-specific. Edges from transcription factors other than TOX are retained where Pando inferred contributions to the same CRE modules; these edges represent broader combinatorial regulatory context and do not designate those factors as TOX cofactors. The lower Venn diagram shows overlap before directional filtering with *TOX*-knockdown bulk RNA-seq: Pando yielded 163 HDF, 185 CTL, and 219 HAEC candidates; 24 were shared across all three cell types and 79 across exactly two. (C) Genome browser view of the *COL25A1* locus, selected as a representative example to examine locus-level support for a Pando-nominated downstream relationship across all three cell types. Chromatin-accessibility tracks and several candidate CREs with peak-to-gene links to the *COL25A1* promoter are shown for HDFs, CTLs, and HAECs across the indicated reprogramming states, and the two representative linked elements selected for sequence-attribution analysis are indicated. (D) State-resolved seq2PRINT sequence-attribution profiles at the two representative linked elements, *COL25A1*-intronic and *COL25A1*-distal. FIMO motif matches to all three Pando-assigned surrogate HMG-box profiles—SOX9 MA0077.1, SOX10 MA0442.2, and TCF7L2 MA0523.1—were detected within each CRE and are displayed as SOX9-like, SOX10-like, and TCF7L2-like matches. The FIMO display maps motif occurrences across the highlighted *COL25A1* intervals and therefore differs from the JASPAR track-labeling format used for the *TOX*-locus panels in Figure 5E and Figure S5B. Pale-yellow blocks delimit the displayed sequence intervals, and horizontal line segments mark the FIMO-predicted match spans. (E) Representative growth/survival-associated signaling modules in HDFs, CTLs, and HAECs. HDFs show a stromal growth-factor/niche signaling module, CTLs show a reprogramming-associated adaptive growth/survival signaling module, and HAECs show a vascular growth/survival signaling module. Linked peak accessibility and gene expression are shown as row-scaled Z-scores. (F) Representative metabolism/transport-associated candidate genes in HDFs, CTLs, and HAECs. Linked peak accessibility and gene expression are shown as row-scaled Z-scores.

